# Cell-death induced immune response and coagulopathy promote cachexia in *Drosophila*

**DOI:** 10.1101/2025.01.07.631515

**Authors:** Ankita Singh, Yanhui Hu, Raphael Fragoso Lopes, Liz Lane, Hilina Woldemichael, Charles Xu, Namrata D. Udeshi, Steven A. Carr, Norbert Perrimon

**Affiliations:** Department of Genetics, Blavatnik Institute, Harvard Medical School, Boston, MA, 7 02115, USA; Broad Institute of MIT and Harvard, Cambridge, Massachusetts, USA; HHMI, Harvard Medical School, Boston, MA, 02115, USA

**Keywords:** Cachexia, Yki^act^ gut tumor, Inflammatory response, DAMPs, Egr/TNFalpha, JNK, Rel, Toll-pathway, PGRP-pathway, Coagulation, Immune response

## Abstract

Tumors can exert a far-reaching influence on the body, triggering systemic responses that contribute to debilitating conditions like cancer cachexia. To characterize the mechanisms underlying tumor-host interactions, we utilized a BioID-based proximity labeling method to identify proteins secreted by Yki^act^ adult *Drosophila* gut tumors into the bloodstream/hemolymph. Among the major proteins identified are coagulation and immune-responsive factors that contribute to the systemic wasting phenotypes associated with Yki^act^ tumors. The effect of innate immunity factors is mediated by NFκB transcription factors Relish, dorsal, and Dif, which in turn upregulate the expression of the cachectic factors Pvf1, Impl2, and Upd3. In addition, Yki^act^ tumors secrete Eiger, a TNF-alpha homolog, which activates the JNK signaling pathway in neighboring non-tumor cells, leading to cell death. The release of damage-associated molecular patterns (DAMPs) from these dying cells presumably amplifies the inflammatory response, exacerbating systemic wasting. Targeting the inflammatory response, the JNK pathway, or the production of cachectic factors could potentially alleviate the debilitating effects of cancer cachexia.

## Introduction

Inter-organ communication is essential for maintaining homeostasis, coordinating physiological processes, and responding to internal and external stimuli. This complex network involves multiple signaling mechanisms, including hormonal, paracrine, and neuronal signaling. Due to these intricate inter-organ interactions, any perturbation in a single tissue can impact the physiology of other tissues. A tumor, for instance, not only affects its immediate microenvironment but also remotely interacts with peripheral tissues through various signaling mechanisms, modulating their metabolic homeostasis ^1–3^. Along with influence on tumor progression and metastasis, these pathophysiological interactions also affect the overall physiological state of the host ^4–9^. One particularly devastating distant tumor-host interaction is cancer cachexia, a metabolic disorder resulting in systemic tissue wasting, severe coagulopathies, increased susceptibility to chemotherapeutic toxicity, heightened immune response, and other lethal sequelae, ultimately leading to reduced quality of life and increased mortality ^10,11^ ^4,11–16^. Despite extensive studies on cancer and its systemic effects, the mediators of cancer cachexia remain largely elusive.

*Drosophila* has been used extensively to study mechanisms of inter-organ communication in both healthy and disease states. In particular, *Drosophila* cancer models have been established to characterize secreted factors from tumors that affect wasting of peripheral tissues ^6,7,17^. A well-established model involves inducing adult gut tumors through the overexpression of an active form of the transcriptional coactivator Yap1 oncogene ortholog Yorkie (yki^3SA^, referred to as Yki^act^) in intestinal stem cells (ISCs). This gut tumor model is associated with metabolic defects and organ wasting phenotypes, such as lipid loss, muscle dysfunction, ovary degeneration, and hyperglycemia ^18–20^. Key proteins secreted by Yki^act^ gut tumors that contribute to organ wasting by disrupting their anabolic/catabolic balance include the insulin-like polypeptide binding protein ImpL2, the Pvr receptor tyrosine kinase ligand Pvf1, and the IL-6-like cytokine Unpaired 3 (Upd3) ^20–22^.

To characterize potential additional factors produced by Yki^act^ gut tumors, we took an unbiased approach to characterize the tumor secretome using the *Escherichia coli* BioID biotin ligase enzyme as a tool for proximity-dependent labeling ^23,24^. Specifically, targeting a promiscuous form of BioID to the secretory pathway in the presence of biotin allows covalent tagging of endogenous proteins within a few nanometers of the labeling enzyme ^25^. Biotinylated secreted proteins are then detected using streptavidin-affinity enrichment, followed by quantitative mass spectrometry (MS). For example, this approach has been used to detect endogenous secreted proteomes in *Drosophila melanogaster, Caenorhabditis elegans*, mouse tumor transplants, and mammalian tissues ^25–28^.

Here, we characterize the secretome of Yki^act^ adult gut tumors, revealing many proteins involved in innate immunity and coagulation, reminiscent to a wound healing response. We demonstrate that the inflammatory response regulates the expression of the cachectic factors *Pvf1, Impl2,* and *upd3*. Additionally, we find that the TNF-alpha homolog Eiger (Egr), a target of Yki in ISCs, activates the JNK and cell death in non-tumorous cells. Our study suggests that targeting the inflammatory response, the JNK pathway, or the production of cachectic factors may provide therapeutic strategies to alleviate the debilitating effects of cancer cachexia.

## Results

### Tumorous guts secrete increased levels of coagulation and immune responsive proteins into the hemolymph

To identify putative cachectic secreted factors from Yki^act^ gut tumors, we expressed endoplasmic reticulum (ER)-targeted BirA* (BirA*ER) in ISCs in control and tumorous guts, and then isolated labeled proteins from hemolymphs using streptavidin (Figure 1f). We used two controls in these experiments. First, as flies with Yki^act^ tumors are produced by crossing the ISC driver *esgGal4, UAS-GFP, tubGal80ts* (EGT) with *UAS-Yki^act^,* one control was generated by crossing EGT with *w*^1118^. Second, as a control for proliferation, we used flies with Ras^act^ tumors, generated by crossing EGT with *UAS-Ras1A*, since expression of the *Ras1* oncogene in ISCs is associated with gut overproliferation with mild organ wasting phenotypes ^9^, compared to flies with Yki^act^ tumors that exhibit severe wasting phenotypes (Figures 1a, 1b,1c, 1d, 1e). Note that, as EGT contains Gal80ts, gut tumors are induced in adults after eclosion by shifting flies to 29°C.

**Figure 1:**
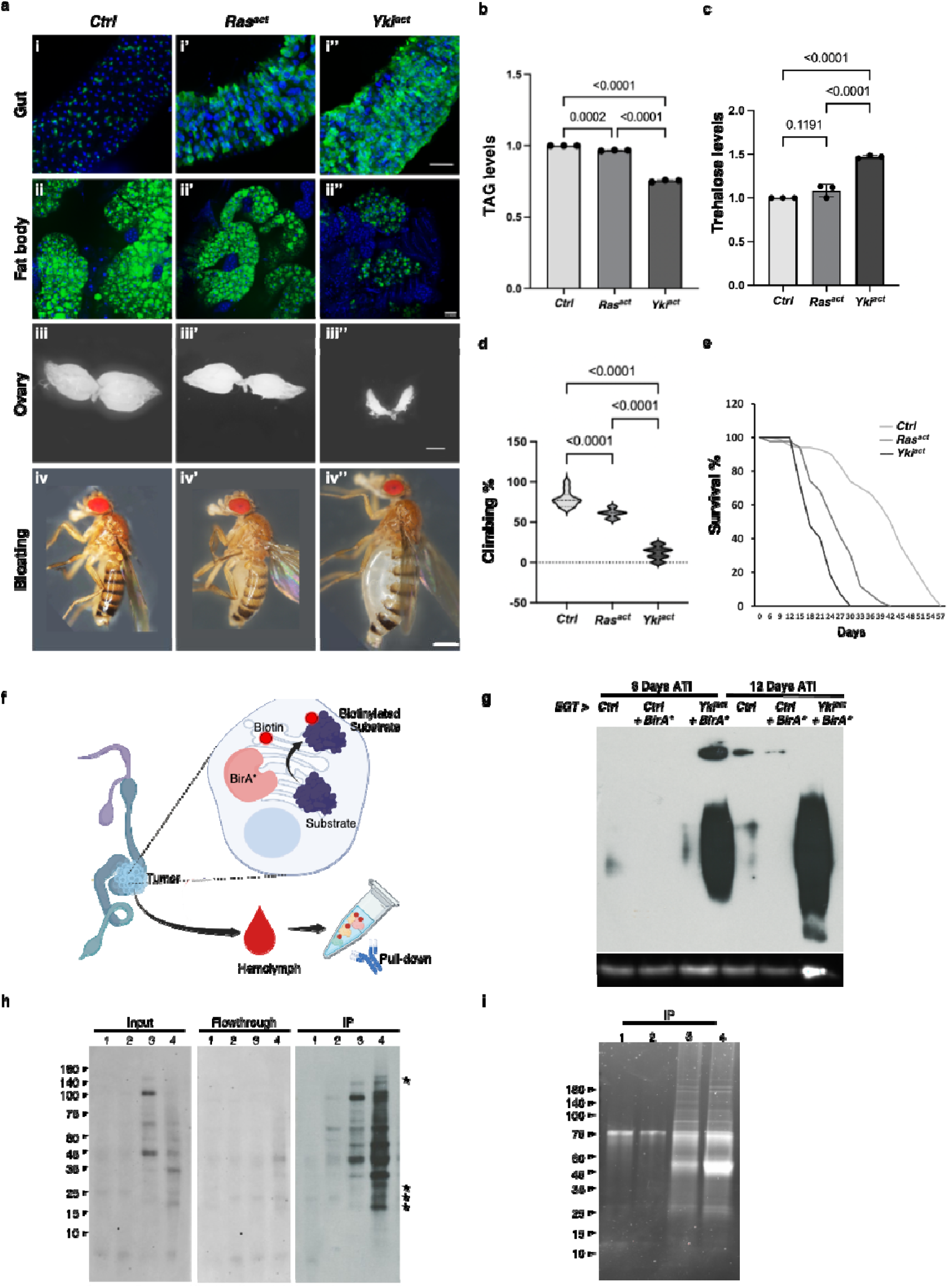
Generation and BirA* labeling of cachectic tumor. (a) Yki^act^ in adult gut ISCs, but not Ras^act^, generates cachectic tumors. First column shows *esgGal4, UAS-GFP, tubGal80ts* driven control flies (*w*^1118^), second column is Ras^act^ flies and third column represents Yki^act^ flies. (i-i’’) Gut tumors, marked by GFP expression, are more pronounced in Yki^act^ flies (i) than in Ras^act^ (i’), and control flies (i’’). GFP labels esg-expressing cells (green) and DAPI stains the nuclei (blue). Scale bar, 10 μm. (ii-ii’’) Bodipy staining (green) of fat body in control and tumor conditions. Adult fat body shows a more pronounced decrease in lipid droplets in Yki^act^ flies than in Ras^act^ and control flies. Scale bar, 10 μm. (iii-iii’’) Brightfield images showing ovary size in control and tumor bearing flies. Yki^act^ tumor causes prominent ovary degeneration. Scale bar, 150 μm. (iv-iv’’). Brightfield images of whole flies showing bloating phenotypes, which was only observed in Yki^act^ flies. Scale bar, 200um. (b) TAG levels of *esgGal4, UAS-GFP, tubGal80ts* adult females (8 females of each genotype in triplicates) after 8 days of tumor induction. (c) Circulating trehalose levels in *esgGal4, UAS-GFP, tubGal80ts* adult females (8 females of each genotype in triplicates) after 8 days of tumor induction. (d) Climbing assay to compare muscle activity of *esgGal4, UAS-GFP, tubGal80ts* flies (n=20). Experiments were performed in triplicates at 12 days. (b, c, d) For statistical analyses, one-way analysis of variance (ANOVA) followed by Tukey multiple test was performed using GraphPad prism software. (e) Lifespan assay to measure viability of *esgGal4, UAS-GFP, tubGal80ts* flies. Y-axis shows percentage of flies that survived, (n=100). Yki^act^ flies have greatly reduced survival compared with control and Ras^act^ flies. (f-i) BirA* labeling of cachectic tumor (f) ER-localized BirA* is expressed in Yki^act^ gut tumors to biotinylate secreted proteins. The promiscuous biotin ligase BirA* biotinylates all ER proteins, which are then secreted into the hemolymph. (g) Expression of BirA* in tumorous Yki^act^ gut tumors labels ER-localized proteins. Immunoblot of adult gut lysates using streptavidin-HRP antibody showing that biotin labeled proteins in Yki^act^ gut tumors are more abundant compared to unlabeled and BirA*-labeled control guts. Anti-Actin was used for internal control. (h) Streptavidin bead immunoprecipitation using hemolymph lysate, followed by streptavidin-HRP immunoblot is used to detect BirA*-labeled proteins secreted in the hemolymph from the gut. The blot on the left is a streptavidin-HRP immunoblot of biotin-labeled proteins from the hemolymph (Input samples). The middle blot shows no remaining labeled proteins in flowthrough after streptavidin (Flowthrough). The blot on the right shows enrichment of biotin labeled proteins after streptavidin-IP (IP). (i) SYPRO Ruby stain of an SDS-protein gel to detect the enrichment of proteins after immunoprecipitation with streptavidin. Samples labeled (h, j). 1: unlabeled control guts; 2: labeled control guts; 3: labeled Ras^act^ gut; 4: labeled Yki^act^ gut. Stars indicate specific bands enriched in hemolymph from the Yki gut.

To ensure proper labeling following BirA*ER expression, we first examined biotin labeling of gut proteins on immunoblots. As expected, labeling was increased in Yki^act^ tumors at 8 and 12 days after tumor induction (ATI) compared to controls (Figure 1g). Next, after confirming the presence of biotin-labeled secreted proteins in the hemolymph of flies at 8 days ATI (Figures 1h, 1i), we performed MS analysis of the biotin-labeled hemolymph proteins using TMTPro 12-plex labeling (Figure 2a, 2b). A total of 3454 proteins (all samples) were identified after MS analysis (Figure 2b, Supplementary Figure 1a). Proteins with an adjusted *P*-value < 0.05 were initially selected and further filtered using a ratio threshold determined as previously described^25^. Using a 10% false positive rate (% FPR) cut-off (Figure 2c), compared to the *w*^1118^ control sample, 162 and 176 candidate proteins were enriched in the Ras^act^ and Yki^act^ samples, respectively (Figure 2e).

**Figure 2.**
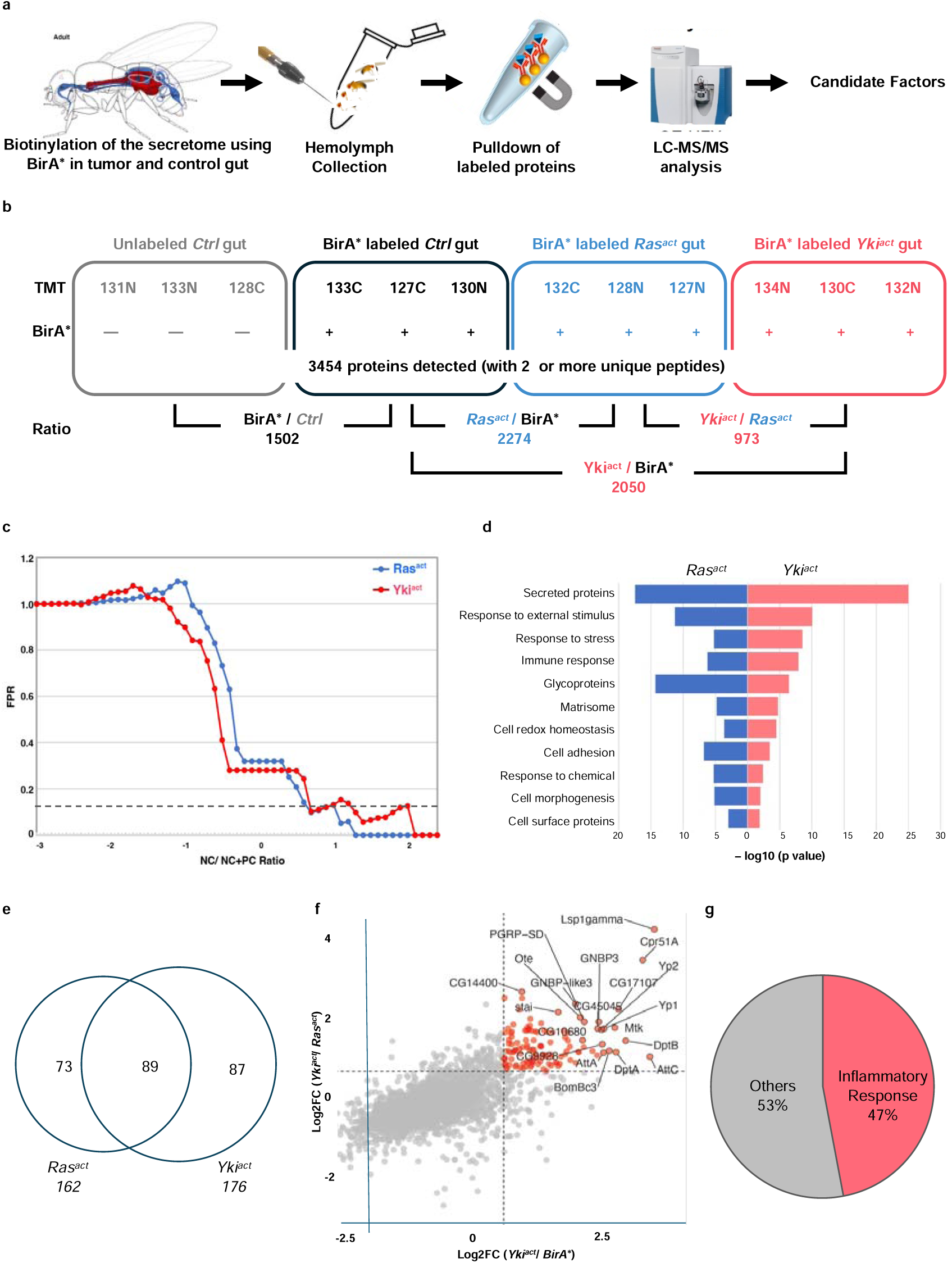
Identification and analysis of Yki^act^ gut tumor-derived proteins in the hemolymph through TMT-MS. (a) Schematic representation of the steps. Biotinylated proteins secreted from Yki^act^ gut tumors are purified from the hemolymph and identified by MS (b) Design of the quantitative proteomic experiment. TMT labels indicate the TMT tags (e.g., 127C) used in all groups. Labeled proteins were immunoprecipitated from fly hemolymph. A total of 12 samples used for analysis-four conditions (Control (Ctrl) gut, BirA* gut, BirA*+Ras^act^ gut, and BirA*+Yki^act^ gut), and three replicates for each condition. (c) Hit identification through NC/NC+PC analysis. X Axis represent NC/NC+PC ratio and Y Axis shows percentage of false positive rate (FPR). The dotted line indicates a FPR threshold < 0.1. Proteins with less than threshold FPR were selected as candidate hits. (d) GO term analysis of the top 100 hits identified in hemolymph of Ras^act^ and Yki^act^ gut tumor bearing flies in comparison to only BirA* gut. The majority of proteins are predicted to be secreted. X-axis represents the negative log10 of p value for enrichment. (e) Overlap plot of hits identified in the Yki^act^ and Ras^act^ secretome in the hemolymph showed over 50% overlap. The numbers in the circle show the number of identified proteins. (f) Scatter plot to show the top 20 labeled hits that were enriched in hemolymph of flies with Yki^act^+BirA* gut tumor compared to Ras^act^+BirA* (Y-axis) and BirA* only (X-axis). The dotted value indicates a Log2FC threshold greater than 1. (g) Pie chart showing that 47% of the top 150 hits belong to inflammatory response proteins.

To identify biological processes associated with the identified proteins, we performed a Gene Set Enrichment Analysis (GSEA) using the PANGEA tool ^29^. The most significantly overrepresented gene group among the hits from both experiments was “secreted proteins” (Figure 2d, Supplementary Figure 1e), which aligns with the expectation that BirA*ER-labeled proteins traffic through the ER. We further analyzed the hits enriched in Yki^act^ samples compared to Ras^act^ samples using gene sets relevant to core biological processes (Supplementary Figure 2). Several known secreted tumor-associated proteins, including Superoxide dismutase (Sod3)^30^, Niemann-Pick type C-2a (Npc2a)^31^, Gelsolin (Gel)^32^, and Nucleobindin1 (NUCB1)^33^, were identified. Further, among the top 100 candidate secreted proteins that were enriched in Yki^act^ compared to Ras^act^ guts, 47 % were associated with inflammatory responses, including cellular response/ coagulation proteins and proteins involved in humoral immunity (Figures 2f, 2g, Supplementary Figure 1f). Finally, to validate our proteomic findings, we analyzed bulk RNA-seq and single-nuclei RNA-seq (snRNAseq) data from Yki^act^ gut tumors ^22,34^ (Table 1; Supplementary Table 1), and observed a strong correlation with our proteomic data, further supporting the biological significance of our findings. Analysis of snRNAseq data revealed that the candidate genes were enriched in ISCs and ECs.

**Table 1:** Top 25 inflammatory response proteins identified in the Yki^act^ secretome. Column 1: Gene name; Column 2: Presence of a signal peptide; Column 3: Association with coagulation or immunity pathways; Column 4: Log fold change in Yki^act^ compared to control (Ctrl); Column 5: Expression enrichment of genes in enterocytes (ECs) of Yki^act^ gut compared to control guts, derived from snRNAseq data^34^. Column 6: Expression enrichment of genes in intestinal stem cells (ISCs) of gut compared to control guts, derived from snRNAseq data; Column 7: Protein size; Column 8: Conservation in humans; Column 9: Reference to their roles in inflammatory pathways.

| Gene | signalP | Pathway | logFC<br>Yki-BioID | ECs<br>(Yki/Ctrl)<br>snRNAseq | ISCs<br>(Yki/Ctrl)<br>snRNAseq | Protein<br>size | Conserved | Reference |
| --- | --- | --- | --- | --- | --- | --- | --- | --- |
| **Lsp1gamma** | Yes | Coagulation | 3.53 | 2.57 | 162.67 | 772 aa | - | <sup>37</sup> |
| **AttC** | yes | Immunity | 3.44 | 12 | 3 | 241 aa | - | <sup>86</sup> |
| **Sumo** | No | Immunity | 3.11 | 2.19 | 4.27 | 90 aa | Yes | <sup>87</sup> |
| **DptB** | yes | Immunity | 2.97 | 7 | 3 | 120 aa | - | <sup>88</sup> |
| **CG17107** | yes | Immunity | 2.83 | 12 | 4 | 96 aa | Yes | <sup>89</sup> |
| **DptA** | yes | Immunity | 2.78 | 6 | 2 | 106 aa | - | <sup>90</sup> |
| **Mtk** | yes | Immunity | 2.76 | 18 | 13 | 52 aa | - | <sup>91</sup> |
| **BomBc3** | yes | Immunity | 2.65 | 12.86 | 31.5 | 98 aa | - | <sup>92</sup> |
| **AttA** | yes | Immunity | 2.54 | 15 | 18 | 221 aa | - | <sup>86</sup> |
| **GNBP3** | yes | Immunity | 2.45 | 2.88 | 4 | 490 aa | yes | <sup>93 94</sup>,<sup>94</sup> |
| **AttB** | yes | Immunity | 2.41 | 6 | 9 | 218 aa | - | <sup>86</sup> |
| **NimB2** | yes | Immunity | 2.32 | 3.38 | 2.8 | 403 aa | Yes | <sup>95</sup> |
| **PGRP-SD** | yes | Immunity | 2.17 | 15.67 | 160 | 186 aa | Yes | <sup>59</sup> |
| **edin** | yes | Immunity | 2.16 | 8.5 | 23 | 115 aa | - | <sup>96</sup> |
| **BaraA1** | yes | Immunity | 2.15 |  |  | 257 aa | - | <sup>97</sup> |
| **GNBP-like3** | yes | Immunity | 2 | 6 | 31 | 152 aa | yes | <sup>94</sup> |
| **PPO2** | No | Coagulation | 1.85 | 3 | 0 | 684 aa | - | <sup>37</sup> |
| **BomT2** | yes | Immunity | 1.45 | 5.44 | 9.83 | 76 aa | - | <sup>98</sup> |
| **MtnA** | No | Immunity | 1.45 | 3.68 | 6.34 | 43 aa | - | <sup>99</sup> |
| **fon** | Yes | Coagulation | 1.3 | 7.67 | 15.67 | 574 aa | - | <sup>38</sup> |
| **PGRP-LB** | Yes | Immunity | 1.25 | 2.97 | 24.73 | 215 aa | Yes | <sup>100</sup> |
| **Idgf5** | Yes | Coagulation | 0.98 | 3 | 8 | 444 aa | Yes | <sup>36</sup> |
| **Idgf6** | Yes | Coagulation | 0.63 | 1.75 | 3.76 | 452 aa | Yes | <sup>36</sup> |
| **Hml** | Yes | Coagulation | 0.39 | 7 | 4.25 | 3842 aa | Yes | <sup>37</sup> |
| **PGRP-SB1** | yes | Immunity | 1.17 | 3 | 4 | 190 aa | Yes | <sup>101 102</sup>,<sup>102</sup> |

### Yki^act^ gut tumors induce a wound healing response

Among the secreted factors identified in our proteomic analysis are many proteins involved in coagulation. These include larval Serum Protein (Lsp1γ), Hemolectin (Hml), fondue (fon), pro-Phenoloxidases (PPOs), Ecdysone-induced gene 71Ee (Eig71Ee), Imaginal Disc Growth Factor 3 (Idgf3), and Tiggrin (Tig). We confirmed the upregulation of these factors in previous bulk RNA-seq and snRNAseq studies (Table 1, Supplementary Table 1), as well as by performing qRT-PCR analysis (Figure 3a).

**Figure 3:**
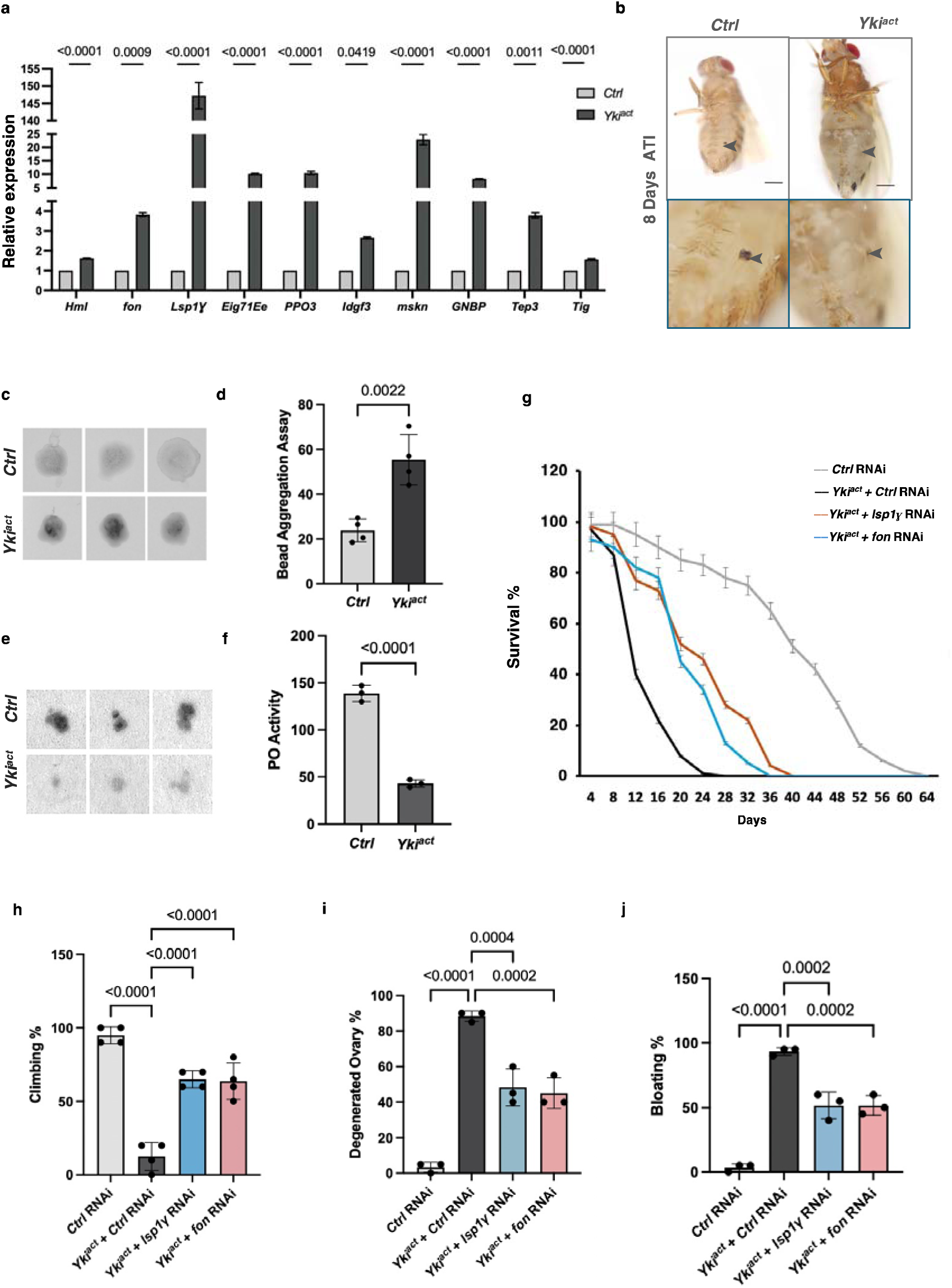
Yki^act^ induced coagulopathy regulates cachexia. (a) qPCR analysis shows that coagulation pathway genes are significantly upregulated in Yki^act^ gut tumors (black bars) compared to control guts (grey bars). Gene expression was normalized to *RP49* and experiments were done in triplicates. Statistical analysis was done using multiple unpaired t-tests. (b) Images of cuticular wounds reveal less melanization in flies with Yki^act^ gut tumors compared to controls (arrowhead marks the wound site). Lower panel shows higher magnification of wound sites. (c) Soft clot activity measurement: Bead aggregation assay showing that hemolymph from Yki^act^ flies with gut tumors has a higher tendency to form aggregates compared to hemolymph from control flies at day 8 ATI (after tumor induction). (d) Bar graph measuring the intensities of the aggregates normalized to Ringer’s solution is shown on the left (n=4). Statistical analysis was done using unpaired t-test. (e) Hard clot activity measurement *in vitro*: PO activity measurement via colorimetry assay using L-DOPA blot reactivity. Hemolymph from Yki^act^ tumor-bearing flies showed reduced L-DOPA blot reactivity compared to control flies (n=9). (f) The graph quantifies the intensity per unit area normalized to background. Statistical analysis was done using unpaired t-test. (g) Early mortality induced in flies with Yki^act^ gut tumors is significantly reduced upon *fon* and *Lsp1*C depletion (n values: = 50) compared to control RNAi depletion in flies with Yki^act^ gut tumors. (h-j) Systemic organ wasting phenotypes observed with reference to climbing activity (n=60) (h), ovary degeneration (n=20) (i), and bloating phenotypes (n=50) (j), were significantly reduced upon knockdown of *fon* and *Lsp1*C. Statistical analysis was done using one-way ANOVA analysis followed by tukey test.

In flies, the coagulation pathway forms an insoluble matrix in the hemolymph to stop bleeding, heal wounds, and protects against infection. Fon, Lsp1γ, and Hml, play a key role in forming the initial “soft clot” ^35–39^, while phenoloxidase (PO)-driven melanization^40–44^ contributes to hardening of the clot “hard clot”. To assess the contribution of coagulation to the phenotypes associated with Yki^act^ gut tumors, we analyzed soft and hard clot formation in these flies. Using a Dynabead aggregation assay, we measured the capacity of hemolymph from tumor-bearing adult flies to form soft clots by inducing clumping of inert Dynabeads (Figures 3c, 3d). Hemolymph from flies with Yki^act^ tumors aggregated beads more readily, indicating heightened soft clotting activity. In addition, Phenoloxidase (PO) activity level in the hemolymph of Yki^act^ flies, measured ex vivo on a colorimetric substrate, was reduced, indicating decreased hard clotting capacity (Figures 3e, 3f). Additionally, flies with Yki^act^ tumors exhibited impaired melanization of both thoracic and abdominal cuticular wounds, and healed more slowly compared to control flies, corroborating the coagulation defects (Figure 3b). Altogether, Yki^act^ tumors overexpress components of the clotting cascade and have coagulation defects. These findings support the hypothesis that tumors are wounds that never heal ^45^, and are consistent with a previous study in *Drosophila* using a different tumor model ^46^.

### Yki^act^ gut tumors induce an inflammatory response

Strikingly, ∼47% of the top 100 secreted proteins identified in the Yki^act^ secretome compared to Ras^act^ gut tumors are associated with innate immunity (Figures 2f, 2g), such as such as Lsp1gamma, PGRPs, GNBPs, Diptericins, Attacins, Drosocins, BomBcs, etc. This enrichment was consistent with previous bulk RNA-seq and snRNAseq analyses ^22^ (Table 1, Supplementary Table 1), which we further confirmed by qRT-PCR (Figure 4a). As innate immunity factors regulate the activity of NFκB transcription factors by nuclear translocation, we examined the subcellular localization of the IMD component Relish (Rel). While Rel was localized in the cytoplasm in control, it was present in the nuclei of Yki^act^ tumorous gut cells (Figure 4d), consistent with activation of the IMD pathway. Interestingly, we also observed an increase in the expression of specific immune-responsive genes, such as *Attacins*, *BomBc*, *Diptericin*, *Drosocin*, *edin*, and GNBP in the fat tissue of flies with Yki^act^ gut tumors compared to control flies, suggesting a systemic immune response to tumors (Figure 4b).

**Figure 4:**
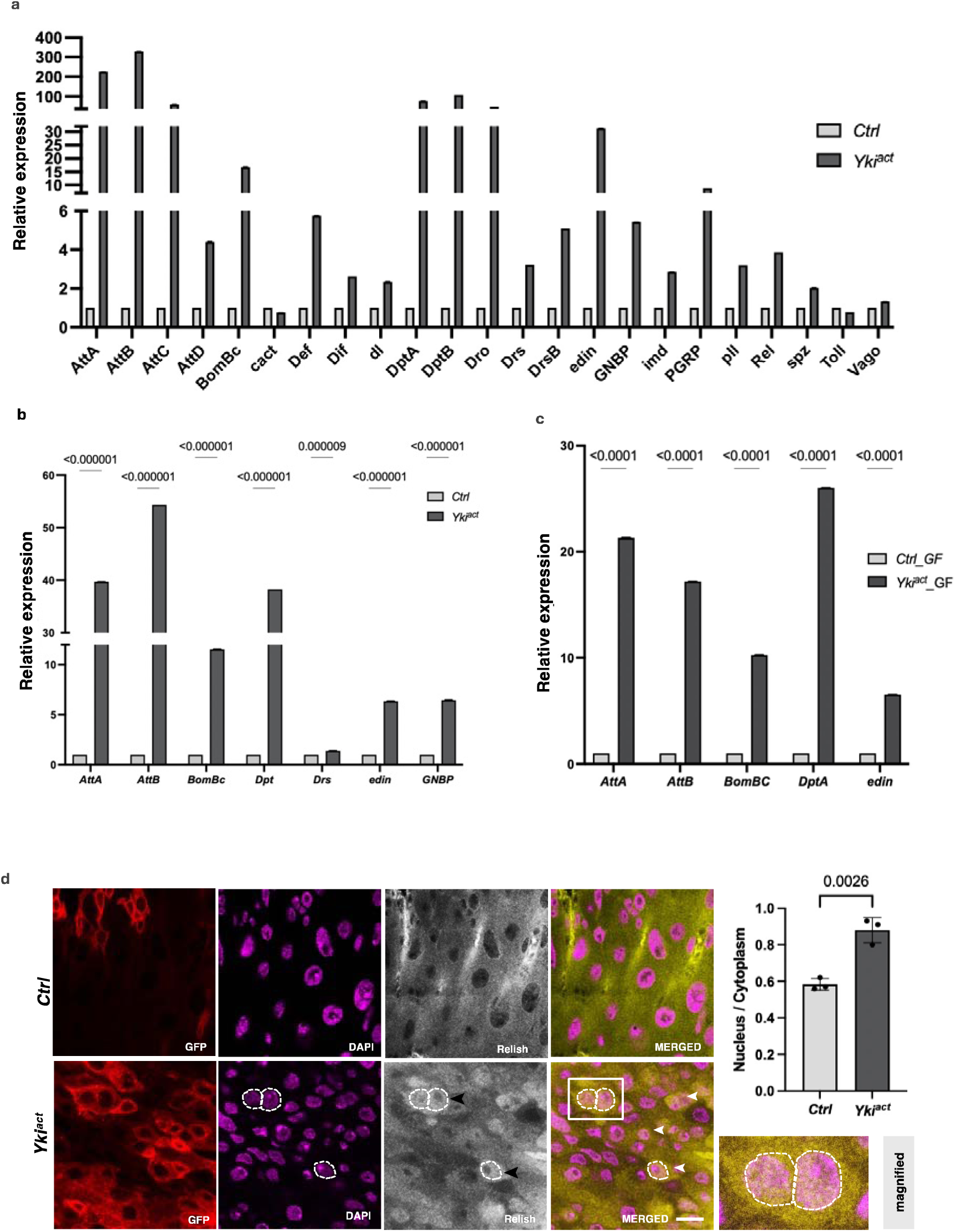
Yki^act^ gut tumors induce innate immune response. (a-c) qPCR analysis shows that immune-responsive genes are significantly upregulated in: (a) Yki^act^ gut tumors (black bars) compared to controls (grey bars), (b) In fat tissue from flies with Yki^act^ gut tumors (dark bars) compared to controls (light bars), axenic (germ-free) compared to control guts (grey bars). Gene expression was normalized to *RP49* and experiments were performed in triplicates. *Attacins* (*AttA, B, C, D*)*, BomBc, Cactus, Defecin* (*Def*)*, Dorsal related immunity factor* (*Dif*)*, dorsal* (*dl*)*, Diptericins* (*Dpt A, B*)*, Drosocin, Drosomycin, edin, metchnikowin* (*met*)*, Gram-negative bacteria binding protein* (*GNBP*)*, imd, PGRPs, pelle (pll*)*, Relish* (*Rel*)*, spatzel* (*spz*)*, Thioester-containing protein 3* (*Tep3*), and *Vago*. (d) Immunostaining for Rel in control and Yki^act^ gut tumors shows nuclear localization of Rel in tumorous guts. False color images are shown here for better contrast. Red indicates *esg*-expressing GFP positive cells. Magenta indicates DAPI staining nucleus, grey indicates Rel localization which is indicated by yellow in merged picture. Nuclear localization of Rel can be seen as orange color due to overlap of yellow (Rel) and magenta (DAPI), indicated by dashed lines and arrowhead. Inset shows the magnified image of boxed area. Scale bar -10um. The nuclear-to-cytoplasmic intensity ratio is quantified in the adjacent bar graph (*n* = 20 nuclei, triplicate readings). Unpaired t-test was used for statistical analysis using GraphPad prism software.

Next, as the induction of innate immunity genes could reflect the presence of bacteria in the gut, we repeated the qRT-PCR experiments in the gut of Yki^act^ and control flies raised in axenic conditions. Increased expression of immune-responsive pathway genes in axenic flies with Yki^act^ gut tumors compared to controls, was still observed, indicating that the tumor itself is sufficient to activate the innate immune response (Figure 4c). However, this increase was not as prominent in axenic flies with tumors as observed in non-axenic flies, suggesting a contribution of the gut microbiota to the overall immune response observed in Yki^act^ tumors.

### Elevated systemic inflammatory response mediates cachexia-like phenotypes in Yki^act^ tumor bearing flies

To explore the contribution of coagulopathy and innate immunity genes to the systemic wasting and early death of flies with Yki^act^ gut tumors, we used RNAi to knockdown genes essential for clotting (*fon, Lsp1g, Tg, Fbp1,* and *Hml*), as well as genes involved in immunity (*PGRPs* and *Toll* receptors, and the NFκB transcription factors *Relish* (*Rel*)*, Dif,* and *dorsal* (*dl*)). Knockdown of *Lsp1*γ and *fon* in Yki^act^ tumor bearing flies significantly rescued systemic effects, and showed increased longevity, reduced bloating, lesser ovary degeneration, and improved muscle activity as measured by climbing assays (Figures 3g-3j). Similar rescued phenotypes were observed with knockdown of *PGRPs, Toll*, *Rel, Dif,* and *dl* (Figures 5a, 5c-5e).

**Figure 5:**
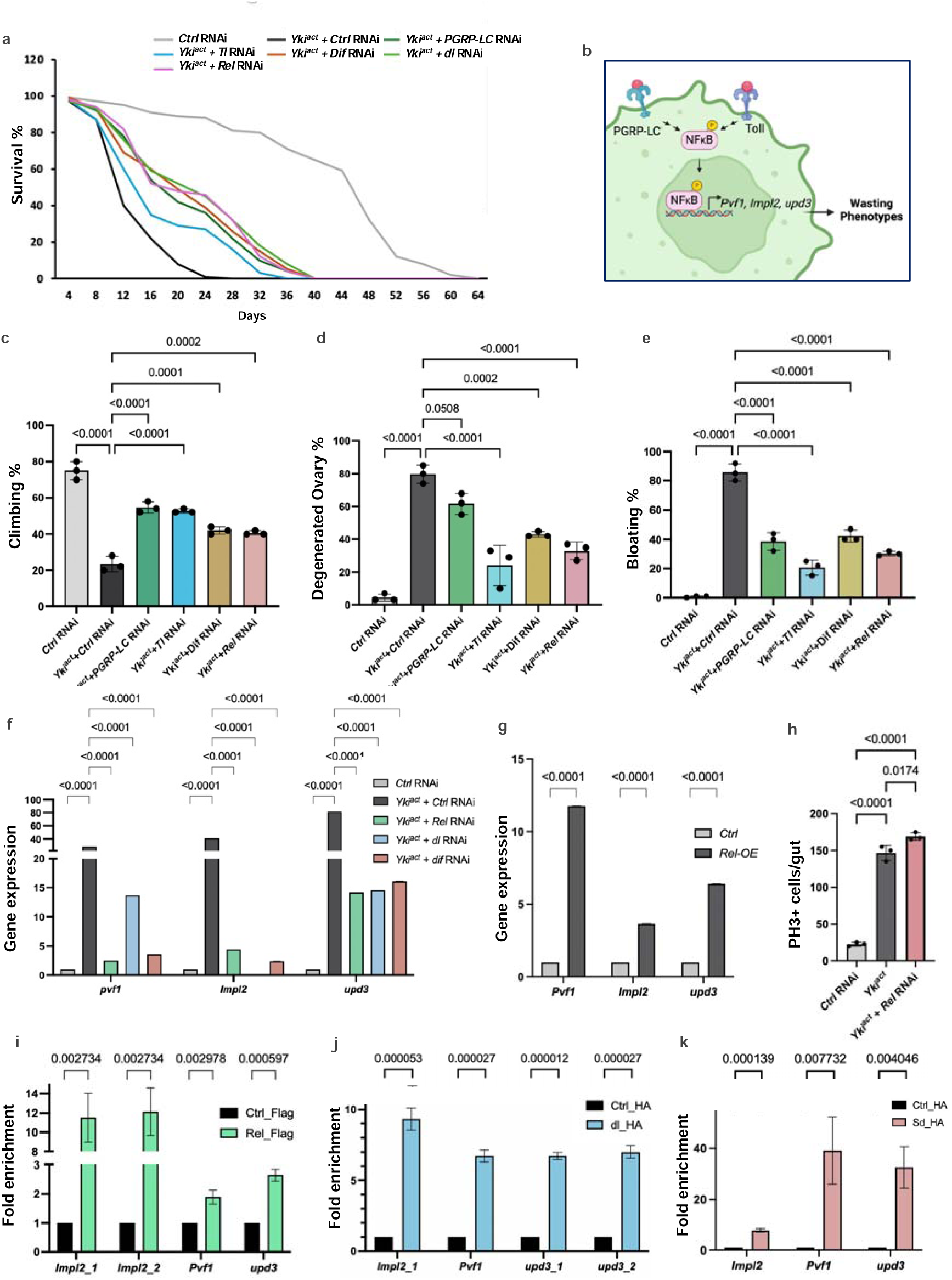
Yki-induced organ wasting phenotypes is mediated by NFκB activation and NFκB transcription factors bind to the regulatory sequences to positively regulate expression of *Pvf1, Impl2* and *upd3*. (a) Early mortality induced in flies with Yki^act^ gut tumors is significantly reduced upon depletion of immunity pathway receptors (*PGRP-LC* and *Toll*) and transcription factors (*Re)*, *Dif*, and *dl*), with *n* = 100 per group, compared to control RNAi depletion in flies with Yki^act^ gut tumors. (b) Schematic to show that NFkB transcription factors can bind to the regulatory sequences of wasting regulating genes-*Pvf1, Impl2 and upd3*. (c) - (e) Systemic organ wasting phenotypes, observed with reference to climbing activity, n= 20 (c), ovary degeneration, n=20 (d), and bloating, n=20 (e), were partially rescued upon knockdown of immunity pathway receptors, *PGRP-LC* and *Toll,* and depletion of transcription factors *Rel* and *Dif.* Three biological replicates were used for the experiment. (f) qPCR analysis showing that knockdowns of *Rel, dl* and *Dif* lead to decreased expression of cachectic marker genes *Pvf1*, *Impl2*, and *upd3.* Relative expression levels are normalized to the *RP49* gene. All experiments were conducted in triplicates. (g) Overexpression of *Relish* increases the expression levels of *Pvf1*, *Impl2*, and *upd3* as seen by qPCR. Three biological replicates were used for the experiment. (h) Graph showing PH3+ cells count upon *Rel* depletion in Yki^act^ guts. To test for significance among the dataset, (c-e, h) one-way analysis of variance (ANOVA) followed by Tukey’s multiple test, and (f, g) a two-way analysis of variance (ANOVA) followed by Tukey’s multiple comparisons test was performed using GraphPad prism software. (i) ChIP assay to show binding of Rel to the regulatory regions of *Pvf1, Impl2* and *upd3* after qPCR using chromatin isolated from *Tubulin-GAL4, Tubulin-GAL80ts>Flag-tagged Relish-68*, compared with the result of the same regulatory regions from non-tagged Ctrl. Immunoprecipitation was done using Flag-beads followed by qPCR. (j) Graph showing the enrichment of *Pvf1, Impl2* and *upd3* regulatory sequences after qPCR using chromatin isolated from *Tubulin-GAL4, Tubulin-GAL80ts>HA-tagged dorsal,* compared with the result of the same regulatory regions from non-tagged Ctrl. Immunoprecipitation was done using HA-beads followed by qPCR. (k) Graph showing the enrichment of regulatory regions of *Pvf1, Impl2* and *upd3* after qPCR using chromatin isolated from *Tubulin-GAL4, Tubulin-GAL80ts> HA-tagged scalloped* (*Sd*), compared to control. qRT-PCR using chromatin isolated from Tubulin-GAL4, Tubulin-GAL80ts> HA-tagged scalloped (Sd) and Ctrl was performed. Immunoprecipitation was done using HA-beads followed by qPCR. For ChIP experiments, three biological replicates were used for the experiments and significance among the dataset was calculated by multiple t-test comparisons using GraphPad prism software. Fold enrichment method was used for normalization.

One explanation for the reduction in systemic wasting observed upon loss of the inflammatory pathway could be an effect on tumor proliferation in Yki^act^ guts. To test, this possibility, we quantified phospho-Histone 3 (PH3)-positive cells in Yki^act^ gut after knock-down of *Rel* compared to in Yki^act^ guts. No reduction in PH3 cell numbers was observed indicating that the suppression of cachectic phenotypes cannot be attributed to a reduction in tumor growth (Figure 5h). Another potential mechanism by which inflammation might exacerbate tumor-associated wasting phenotypes is through the regulation of the expression of cachectic factors in tumor cells. Thus, we examined whether the NFκB transcription factors, Rel, dl and Dif, could regulate directly the expression of *Pvf1, Impl2,* and *upd3,* previously identified as factors secreted from Yki^act^ gut tumors that trigger wasting ^20–22^. Peak annotations of Chromatin immunoprecipitation (ChIP)-seq datasets for Rel and dl/Dif from the modERN website^47^ revealed that they can bind near the transcription start site (TSS) of *Pvf1, Impl2*, and *upd3* (Supplementary Figures 3a, 4a-4c), therefore we considered them as the potential regulatory sequences. Furthermore, Alphafold 3 tool, that designs the 3D structure of proteins and model protein–DNA interactions, also supports that these *Pvf1, Impl2,* and *upd3* regulatory sequences can bind to Rel and dl/Dif with strong iPTM scores (Supplementary Figure 3b). To validate these observations, we performed ChIP of expressed Flag-tagged cleaved Rel (Flag-Rel), HA-tagged dorsal (HA-dl) and control flies, using anti-Flag and anti-HA magnetic bead respectively, followed by quantitative PCR (ChIP-qPCR) using primers specific to the regulatory sequences of *Pvf1, Impl2,* and *upd3*. ChIP-qPCR analysis revealed an enrichment of *Pvf1, Impl2,* and *upd3* regulatory sequences in chromatin samples expressing FLAG-Rel and HA-dorsal, compared to the control chromatin samples (Figures 5i, 5j). Further, consistent with our results that NFκB transcription factors can bind to the regulatory sequences of the cachectic factors, overexpression of *Rel* in wild-type guts was sufficient to increase the expression of *Pvf1, Impl2,* and *upd3* (Figure 5g). In addition, knockdown of *Rel, dl,* and *Dif* in Yki^act^ gut tumors reduced the expression of *Pvf1, Impl2,* and *upd3* (Figure 5f). Altogether, these results indicate that Yki^act^ gut tumors trigger an immune response that positively regulates the expression of the cachectic factors.

We further examined whether overexpression of *Rel* in wild-type ISCs itself is sufficient to induce tumors or cachexia, but we did not observe any significant overgrowth or cachexia phenotypes indicating that there are additional factors that cooperate with Rel to induce cachexia in Yki^act^ tumor bearing flies. Finally, as removal of *PGRPs, Toll*, *Rel, Dif,* and *dl* (Figures 5a, 5c-5e) from Yki tumors only partially suppress *Pvf1, upd3* and *Impl2*, we wondered whether Yki/Sd itself could also regulate the expression of these cachectic factors. ChIP-qPCR analysis on flies expressing HA-tagged Sd (HA-Sd) and control flies, using primers specific to the regulatory sequences of *Pvf1*, *Impl2*, and *upd3*, demonstrated a significant enrichment of these regulatory sequences in chromatin samples expressing HA-Sd compared to controls (Figures 5k). Altogether, these findings suggest that in Yki^act^ guts, the expression of cachectic factors is co-regulated by the Yki/Sd transcription complex and inflammatory transcription factors.

### Yki^act^ gut tumors activate Eiger/JNK signaling to induce cell death in neighboring cells

Cell death induced during tumor growth or cell injury can cause inflammatory responses via the release of DAMPs (damage-associated molecular patterns)/CDAMPs (cell death-associated molecular patterns) ^48,49^ Thus, we examined the extent of cell death/apoptosis in flies with Yki^act^ gut tumors as a possible trigger of the innate immune response. Immunostaining for the cleaved (activated) apoptosis marker DCP1 (Death Caspase-1/active caspase-3) revealed a significant increase in apoptotic cells in Yki^act^ gut tumors compared to both control and Ras^act^ tumor-bearing flies (Figures 6a, 6d). Interestingly, DCP1 positive cells were GFP negative (GFP marks Yki^act^ expression), indicating that Yki^act^ induction causes cell death in neighboring GFP negative wild-type cells in a cell non-autonomous manner (Figure 6aiii). Presumably, DAMPs are released by these dying cells, as in Yki^act^ guts, compared to Ras^act^ (data not shown) and control guts, an increase in reactive oxygen species (ROS) levels, a key component of DAMPs ^50^, was observed. This increase was detected using Dihydroethidium staining (Figures 6b, 6e). Released DAMPs from dying cells probably trigger the inflammatory response observed in Yki^act^ guts.

**Figure 6:**
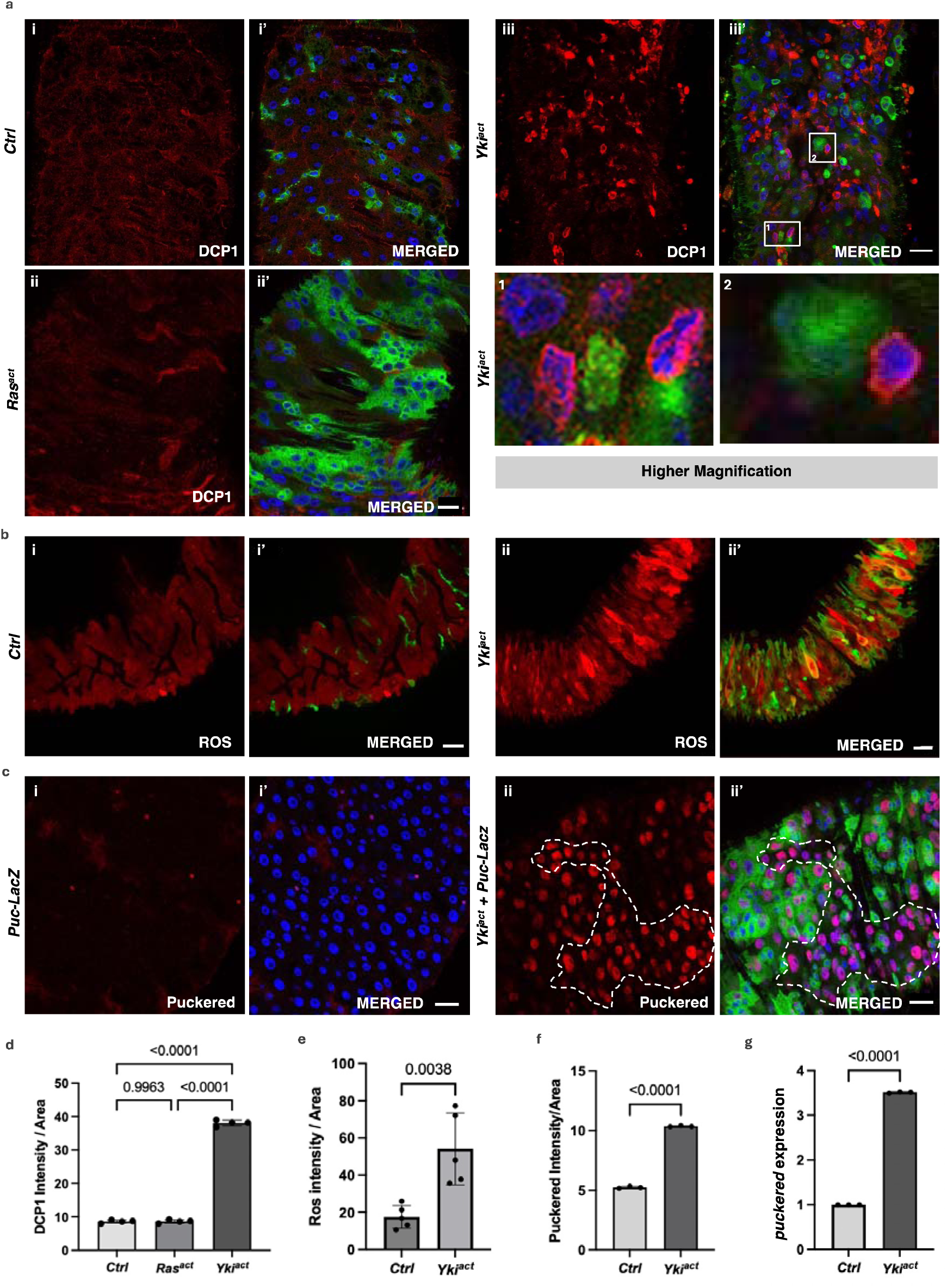
Yki^act^ gut tumors induce JNK-mediated cell death in neighboring non-tumor cells. (a - c) Immunostaining images where GFP marks *esg*-positive cells; blue shows DAPI-stained nuclei; and red is for specific staining. (a) DCP1 staining to assess cell death in the guts of (i, i’) control, (ii, ii’) Ras^act^, and (iii, iii’) Yki^act^ flies. The first column displays DCP1 staining in red (i, ii, iii); and second column is the merged image (i’, ii’, iii’). Extensive cell death was observed in Yki^act^ gut compared to Ctrl and Ras^act^. Scale bar, 20 µm. 1 and 2 images are inset images of boxed area 1 and 2 respectively, showing cell-death in neighboring cells of Yki^act^ GFP+ cells. (b) Dihydroethidium (DHE) dye staining to assess ROS production in the guts of (i, i’) control, and (ii, ii’) Yki^act^ guts. The first column displays ROS staining in red (i, ii); and second column is the merged image (i’, ii’). Heightened ROS production was observed in Yki^act^ guts confirming cell damage. Scale bar, 20 µm. (c) β-Galactosidase staining to mark the expression of *puckered-LacZ (puc-lacZ)*, a known reporter of JNK activity, is shown in red. Increased *puc* expression was observed in Yki^act^ guts compared to Ctrl guts, specifically, non-GFP cells showed higher *puc* expression compared to GFP-positive Yki^act^ expressing cells. Scale bar, 20 µm. Dashed lines mark the area of higher puckered expressing non-GFP cells. (d) - (f) The graph represents quantification of the average intensity of DCP1 staining per unit area (d), ROS per unit area (e), and *puc* staining per unit area (f), normalized to background, using ImageJ. Light colored bar represents Ctrl while dark colored bar represents Yki^act^ guts. Three or more images were used for quantification of intensities. To test for significance among the datasets, a one-way analysis of variance (ANOVA) (d) or unpaired t test (e, f) followed by Tukey’s multiple comparisons test was performed using GraphPad prism software. (g) qPCR analysis to show increased expression of *puc* in Yki^act^ guts (dark bar) compared to Ctrl (light). Expression levels were normalized to *RP49*, and experiments were conducted in triplicates. To test for significance among the datasets, unpaired t test followed by Tukey’s multiple comparisons test was performed.

One possible mechanism by which Yki^act^-expressing cells might cause non-autonomous cell death in neighboring cells could be by inducing cell competition through JUN kinase (JNK) signaling ^51,52^. Thus, we examined the expression of *puckered* (*puc*)-*LacZ*, a known reporter of the JNK signaling pathway, in Yki^act^, Ras^act^ and control guts. *puc* expression was massively upregulated in Yki^act^ gut tumors compared to Ras^act^ (data not shown) and controls (Figures 6c, 6f), a result consistent with mRNA expression detected by qPCR (Figure 6f). Next, as JNK signaling is known to be activated by the TNF-alpha ligand Eiger (Egr) ^53,54^, we examined the expression of *egr* in Yki^act^ gut tumors using qPCR which was increased compared to control (Figure 7a), a result consistent with snRNAseq data^34^ (Supplementary figure 5c).

**Figure 7:**
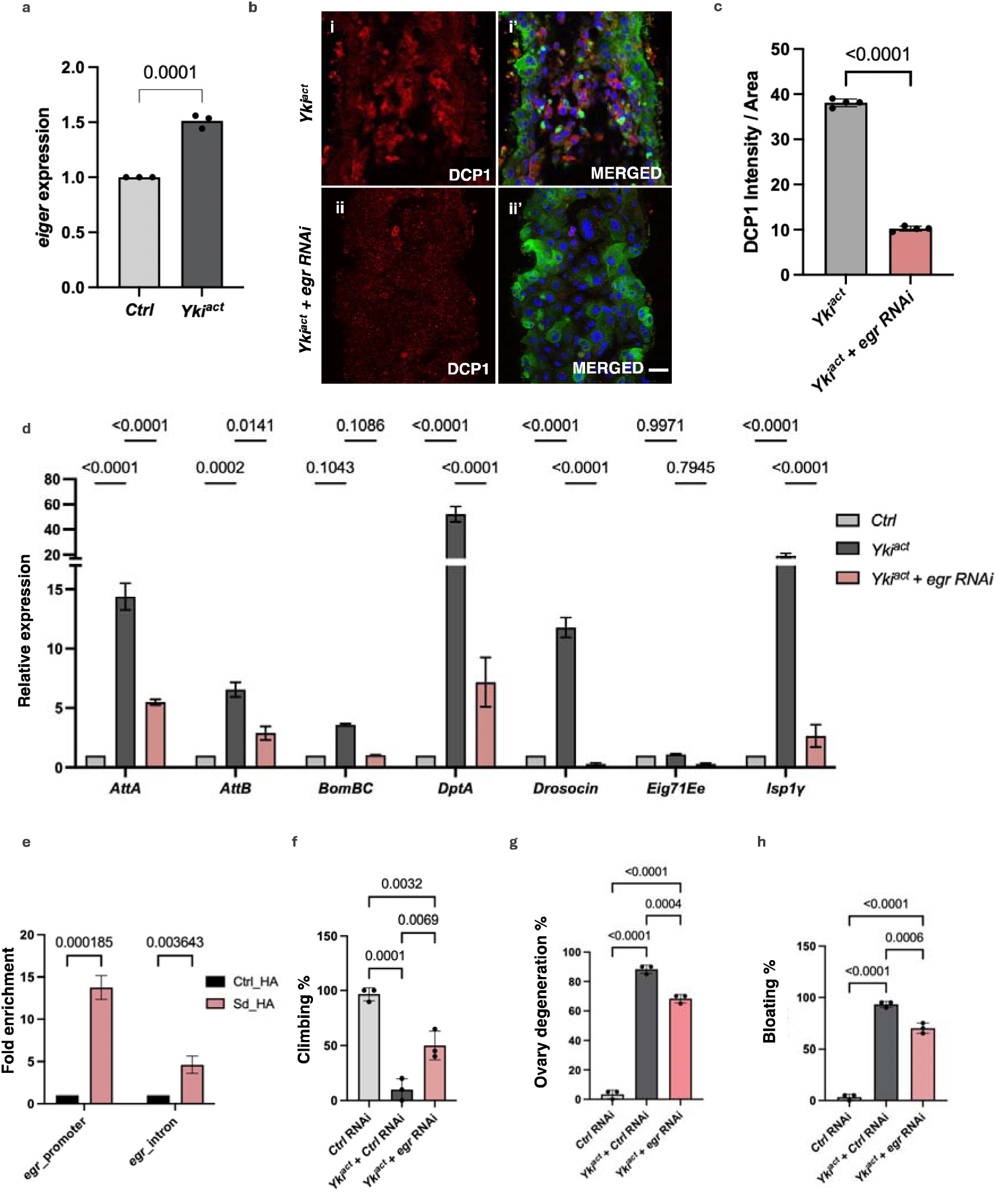
Regulation of cell death by Eiger. (a) qPCR analysis demonstrates elevated expression of *eiger (egr)* in guts (black bars) compared to control guts (grey bars). Experiment was conducted in triplicates. To test for significance among the dataset, unpaired t test was performed using GraphPad prism software. (b) DCP1 staining to assess cell death in the guts of (i, i’) *Yki^act^*, and (ii, ii’) *Yki^act^* + *egr-RNAi* guts. The first column displays DCP1 staining in red (i, ii); and second column is the merged image (i’, ii’). Extensive cell death was observed in Yki^act^ gut which was rescued upon knock-down of *egr*. Scale bar, 20 µm. (c) Quantification of the average intensity of DCP1 staining per unit area normalized to background, using ImageJ. Dark colored bar represents Yki^act^ while peach colored bar represents *Yki^act^ + egr RNAi* guts. Three or more images were used for quantification of intensities. To test for significance among the datasets, unpaired t test was performed. (d) Rescue of inflammatory gene expression by knockdown of *egr*. Relative expression of inflammatory response genes is reduced in *Yki^act^* + *egr-RNAi* samples compared to *Yki^act^*guts. Gene expression was normalized to *RP49*. Experiments were conducted in triplicates. Significance among the dataset was calculated by a two-way analysis of variance (ANOVA) followed by Tukey’s multiple comparisons test using GraphPad prism software. (e) ChIP assay to show binding of Scalloped (Sd) to regulatory regions of *egr* after qPCR using chromatin isolated from *Tubulin-GAL4, Tubulin-GAL80ts> HA-tagged scalloped* (Sd), compared to the result of the same regulatory regions from non-tagged Ctrl. Immunoprecipitation was done using HA-beads followed by qPCR. Fold enrichment method was used for normalization. For ChIP experiments, three biological replicates were used for the experiments and significance among the dataset was calculated by a two-way analysis of variance (ANOVA) followed by Tukey’s multiple comparisons test using GraphPad prism software. (f-h) Systemic organ wasting phenotypes observed with reference to climbing activity (n=60), ovary degeneration (n=20) and bloating phenotypes (n=50), were significantly reduced upon knockdown of *egr*. Statistical analysis was done using one-way ANOVA analysis followed by tukey test.

As Yki associates with the transcription factor Scalloped (Sd) in the nucleus to promote downstream target gene expression ^55^, *egr* expression may be directly regulated by Yki/Sd. Peak analysis of ChIP-seq datasets of Sd from the modERN website^47^ revealed Sd binding near the TSS of *egr* (Supplementary Figures 5a, 5d). Further, Alphafold 3 supported that these regulatory sequences of *egr* might bind to Sd with a strong iPTM score of 0.9 (Supplementary Figure 5b). To directly test whether *egr* is a transcriptional target of Yki/Sd, we performed ChIP-qPCR of expressed HA-tagged Sd (HA-Sd) and control flies using primers specific to the regulatory sequences of *egr*. *egr* regulatory sequences of chromatin samples expressing HA-Sd were significantly enriched (Figure 7e), confirming that Yki/Sd can bind to the promoter sequences of *egr* to regulate its expression.

Finally, to test the role of *egr* in Yki^act^ mediated cell death, we knocked down *egr* in Yki-expressing cells and checked the extent of cell death and expression of inflammation responsive genes. Immunostaining of cleaved DCP1 revealed a significant rescue in Yki^act^-induced apoptosis upon *egr* knockdown (Figure 7b). Additionally, knockdown of *egr* in Yki^act^ tumors caused a substantial down-regulation of the expression of genes involved in cell death-induced inflammatory responses (Figure 7d) and partially rescued systemic effects like reduced bloating, lesser ovary degeneration, and improved muscle activity (Figure 7f - 7h). Altogether, these results suggest that induction of *egr* in Yki^act^ tumor causes JNK-mediated cell death, ultimately resulting in inflammatory responses.

## Discussion

In this study, we used a proximity labeling approach to identify proteins secreted by *Drosophila* Yki^act^ gut tumors associated with systemic organ wasting. Among the secreted factors enriched in the Yki^act^ tumor secretome, we identified multiple proteins involved in *Drosophila* innate immunity, particularly within the inflammatory and coagulation pathways. Characterization of the regulation of the innate immunity pathway leads us to propose the model that: 1. Yki/Sd regulate the expression of Egr/TNF-alpha in tumorous cells; 2. Egr activates JNK signaling in non-tumorous cells leading to cell death; 3. Release of DAMPS by dying cells activate the innate immunity pathway/NFkb in tumorous cells; and 4. Activation of NFκB transcription factors regulate the expression of innate immunity factors as well as the cachectic factors *Pvf1, Impl2,* and *upd3* (Figure 8).

**Figure 8:**
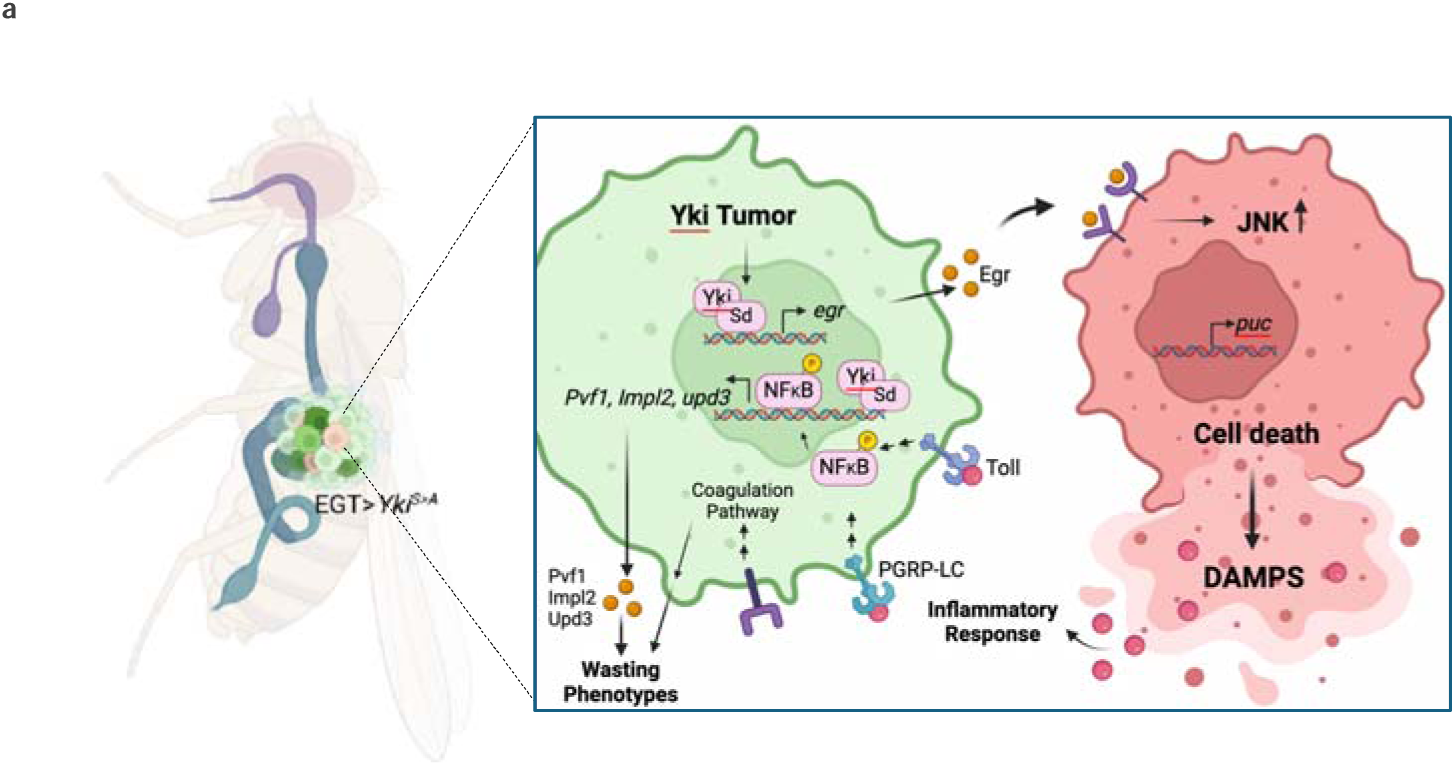
Model. Yki^act^ activation stimulates Egr/JNK signaling, leading to cell-competition-driven cell death in adjacent non-tumor cells. Egr/JNK activity initiates an inflammatory response within the tumor, potentially due to damage-associated molecular patterns (DAMPs) from dying cells. This non-autonomously induced inflammatory response, combined with cell-autonomous Yki/Sd transcription factors, collectively regulates the expression of *Pvf1*, *upd3*, and *Impl2*.

Although our study focuses on proteins secreted by *Drosophila* Yki^act^, we provide a resource of secreted proteins that are enriched in Yki^act^ tumors and Ras^act^ tumors in comparison to controls (Supplementary Table 1). It is noteworthy that we did not identify known secreted ligands from Yki^act^ tumors, including Pvf1, Impl2, and Upd3, which may be due to a number of reasons: 1. lysine residues that are labeled by biotin may not be accessible in some proteins due to their in vivo secondary and tertiary structures; 2. in vivo detection might miss proteins that are part of large complexes, e.g., if they are located in deep pockets of a complex; 3. some proteins might be secreted via unconventional pathways and not labeled by the ER-localized BirA* enzyme; and 4. the MS analysis focused on proteins with more than two distinct peptides, potentially leading to the exclusion of smaller proteins.

*Drosophila* innate immunity comprises two main responses: the humoral and cellular systems. Our analysis revealed that the Yki^act^ secretome is enriched in proteins involved in the cellular response, exemplified by clotting cascade proteins that activate hemocytes, as well as proteins involved in the humoral response via NF-κB-mediated pathways. The soft clot and hard clot forming components abundantly expressed in Yki^act^ secretome suggest that gut tumors engage with the host clotting cascade. Consistent with our findings, a recent report using a *Drosophila* ovarian carcinoma tumor model also found that attenuation of coagulopathy extends lifespan of tumor-bearing hosts ^46^. As in the ovarian model, we observed a reduced tendency to form melanized clots and decreased phenoloxidase (PO) activity. This transition might be attributed to the overstimulation of the clotting system by the tumor, leading to exhaustion of clotting components within the host. For example, crystal cells, which are produced only during the larval stage but persist into adulthood, might become depleted ^56^. A similar phenomenon is observed in certain human cancers and is known as disseminated intravascular coagulation (DIC), a syndrome characterized by widespread intravascular activation of coagulation coupled with the simultaneous consumption of coagulation factors and platelets ^57^. In addition, clotting proteins like Fon, Tig, and Lsp1y have been reported to influence functions beyond coagulation, such as muscle attachment ^58^, suggesting that they might contribute to muscle wasting. Remarkably, loss of either *lsp1*γ or *fon* led to an alleviation of wasting phenotypes, and ultimately an extension of lifespan. These findings suggest that coagulating proteins can influence the Yki^act^ associated wasting phenotypes and early lethality, and align closely with Dvorak’s characterization of tumors as “wounds that never heal”^45,46^.

Strikingly, the most prevalent class of proteins secreted from Yki^act^ tumors are associated with the immune response pathway. *Drosophila* innate immunity is primarily governed by the NF-κB pathways, which are represented by two key recognition and signaling cascades: the Toll and IMD pathways ^59,60^. The observed enrichment of NF-κB pathway proteins in our proteomic data was confirmed through several independent transcriptomic analysis. Collectively, these findings indicate that Yki^act^ tumors activate NF-κB signaling within the gut, a phenomenon that mirrors observations in several human carcinomas where the upregulation of the NF-κB pathway creates a supportive microenvironment critical for tumor initiation and progression ^61^. Furthermore, to address the possibility that the heightened immune response observed in Yki^act^ gut tumors might be due to changes in microbiota composition or damage to the peripodial membrane, which could expose gut cells to microbiota and trigger an immune response, we assessed the expression of key inflammatory genes in axenic flies, which are free of microbiota. qPCR analysis revealed a significant increase in the expression of inflammatory genes in Yki^act^ tumor axenic guts compared to control axenic guts, confirming that Yki^act^ tumor itself is sufficient to induce an increased inflammatory response. However, the increase in inflammatory gene expression was not as pronounced as in non-axenic Yki^act^ flies. This suggests that the gut microbiota can impact the immune response in Yki^act^ gut which is consistent with previous studies reporting that the gut microbial community and its homeostasis can influence the inflammatory response in multiple cancers ^62^

To further investigate whether the upregulation of NF-κB pathway proteins could be considered a “tumor-promoting inflammation” response—a phenomenon commonly observed in human carcinomas—we performed loss-of-function experiments targeting Toll and Imd pathway components. RNAi-mediated knockdown of Toll and Imd pathway components partially rescued Yki^act^-induced wasting phenotypes and premature death, indicating that NF-κB signaling contributes to Yki^act^-induced cachexia. In addition, we demonstrated binding of NFκB transcription factors to *Pvf1, Impl2* and *upd3* regulatory sequences. Our findings that activation of the humoral immune response can lead to wasting phenotypes, is further supported by another study, which demonstrated that flies infected with bacterial pathogen *Mycobacterium marinum* undergo organ wasting ^63^. Although mammalian studies have demonstrated the role of the NF-κB pathway in cancer cachexia-induced skeletal muscle wasting ^64^, our study is the first to show that Yki^act^-induced activation of NF-κB in tumors drives host wasting, including muscle dysfunction, ovary degeneration, and host survivability, through the regulation of cachectic factors. Importantly, this finding provides a mechanistic link between tumor-induced inflammation and cachexia, shedding light on how immune signaling pathways drive tissue wasting during tumor progression. In addition, the coagulopathy observed in the Yki^act^ gut may be linked to NF-κB activation, as previous human studies have reported that inflammatory responses can significantly impact the clotting process by upregulating the expression of pro-coagulant factors ^65,66^.

Importantly, we provide evidence that cell death drives the inflammatory response activated by Yki^act^ tumors. During tumor growth or cell injury, dying or apoptotic cells release various stimuli that can trigger a sterile inflammatory response. Reports from human studies indicate that successful phagocytosis of these dying cells can prevent the activation of the inflammatory response. However, in patients with impaired phagocytic activity, apoptotic cells are not properly processed, leading to a necrotic pathway where cells lose plasma membrane integrity. This results in the release or exposure of intracellular components, including endogenous materials from the cytosol, nucleus, and mitochondria, which are recognized as pro-inflammatory molecules. These molecules, collectively referred to as alarmins, DAMPs, or CDAMPs, are endogenous danger signals released from damaged or dying cells into circulation ^48,49,67^. In circulation, these DAMPs trigger a DAMP-Associated Response (DAR), activating the innate immune system through interaction with pattern recognition receptors (PRRs) or by activating hemocytes to promote wound healing responses^68^. While DAMPs play a role in host defense, they can also contribute to pathological inflammatory responses, as previously shown in studies of tumor microenvironments ^69^. In flies, well-characterized secretory DAMPs include Actin, ATP, Calreticulin, ROS, Egr, and Spz, which might be exposed at injury sites or released into circulation^50^. To confirm DAMP-induced activation of the inflammatory response in Yki^act^ tumors, we first validated increased cell death, marked by DCP1 staining, in Yki^act^-expressing guts compared to Ras^act^ and control guts, and found that dying cells, in turn, activate DAR, as revealed by increased ROS levels in Yki^act^ gut, to initiate an inflammatory response. None of the DCP1-positive cells were Yki^act^ -expressing GFP-positive cells, consistent with previous findings that activation of Yki triggers the transcription of target genes that are well-characterized for either promoting cell proliferation or suppressing apoptosis ^70^. Given that all dying cells were non-tumor cells, we speculate that in the tumorous gut, Yki^act^ induces cell proliferation autonomously while causing cell death non-autonomously in nearby wild-type cells, a phenomenon reminiscent to ’supercompetition’ ^71^. This model is supported by previous reports in other cellular contexts, where loss of Hippo pathway activation components or activation of Yki can transform cells into super-competitors, inducing cell death in neighboring cells ^72,73^. In summary, our study supports the model that Yki^act^ tumor cells non-autonomously induce cell death in neighboring non-tumorous cells, thereby triggering a DAR and initiating an inflammatory response.

Finally, we showed that Yki^act^ induce cell death in non-tumor cells through the regulation of Egr/TNF alpha and /JNK signaling. In *Drosophila*, there is a single TNF ligand, Egr, a potent inducer of apoptosis ^54^, and its pro-apoptotic function depends entirely on its ability to activate the JNK pathway ^53^. Numerous studies have suggested that JNK signaling in *Drosophila* can be both pro-apoptotic and pro-proliferative depending on the cellular context. We observed that the non-autonomous cell death is mediated by Egr-induced JNK signaling, consistent with the established role of JNK in promoting cell competition and tissue homeostasis ^53^ ^74^. Through various genetic and molecular approaches, we demonstrated that the Yki partner Sd can bind to the regulatory sequence of *egr*, to upregulate its expression. Overexpressed Egr is presumably secreted to activate JNK signaling in the gut. Further, expression analysis of *puc-LacZ*, an established marker for JNK activation, revealed two distinct levels of JNK activation in different cell populations of Yki^act^ gut tumors: non-GFP cells showed very high expression (marked by white dotted lines, Figure 6c), while GFP-positive *Yki^act^*^-^expressing cells, primarily ISC/EB cells), exhibited low JNK activation. This differential signaling likely plays a key role in tumor dynamics. Interestingly, it has been reported that low levels of JNK activity in ISCs promote ISC proliferation, while high levels trigger apoptosis ^75^. Thus, considering the pattern of *puc-lacZ* expression in Yki^act^ guts, we speculate that Yki^act^-expressing cells strategically maintain low JNK activity to sustain and promote their proliferation, while simultaneously inducing high JNK activity in neighboring non-tumor cells to cause cell death via non-cell autonomous mechanisms. Furthermore, prior studies in both *Drosophila* and mammals have demonstrated that activated NF-κB transcription factor, Rel, can attenuate JNK activation through the proteasomal degradation of TAK1, a key JNK regulator ^76–78^. In Yki^act^ gut tumor cells, the observed increased in Rel activity could act as a protective mechanism, limiting JNK hyper-activation within tumor cells, and thereby preventing cell death and promoting proliferation. This interplay between Rel and JNK signaling could be a potential mechanism by which Yki^act^ tumors evade apoptosis while inducing inflammation and cell death in the surrounding tissue. As a corollary, we have shown that Yki^act^ can trigger Egr/JNK signaling to induce cell death in non-tumor cells. Interestingly, *egr* expression has been shown to stimulate ISC proliferation following overexpression of Ras in ISC/EB ^79^ or during injury ^80^, or in more physiological conditions in an autocrine and paracrine manner ^81^. Thus, it is possible that Egr in addition to its non-autonomous role in triggering cell death, also acts autonomously to contribute tumor cell proliferation. Finally, Imd-NFκB has been shown to synergize with JNK pathway for enterocytes (ECs) shedding upon infection via a mechanism independent of ROS-associated apoptosis ^82^, suggesting that EC shedding may also play a role in the development of Yki tumors.

The loss of *egr* and inflammatory factors such as *Rel* and *dif* only partially rescued the cachexia phenotypes and the expression of cachectic factors in Yki^act^ tumors, leading us to explore whether Yki, together with its co-factor Sd, directly regulate *pvf1*, *Impl2*, and *upd3* expression. Our findings suggest that, in addition to NFκB factors, Yki/Sd also regulate *pvf1*, *Impl2*, and *upd3*, indicating that the cachectic factors integrate both cell-autonomous and non-cell-autonomous mechanisms (Figure 8). Altogether, our results suggest that direct intervention in inflammatory response pathways or in the production of cachexia-inducing factors offers significant potential as a therapeutic approach to counteract the debilitating effects of cancer cachexia.

## Materials and Methods

### Drosophila stocks

All fly stocks were maintained on a standard cornmeal, yeast, molasses, and agar diet at 25°C. Wild-type control flies used in experiments were *w*^1118^. Crosses were performed at 18°C with the same food conditions. For temperature-sensitive experiments, the initial crosses were set up at 18°C, and the adult flies were transferred to 29°C. *UAS-yki*^*[S3A]*^ (*w*;;UAS-yki.S111A.S168A.S250A.V5;* 228817), which we refer to as *UAS-yki^act^*, was obtained from the Bloomington *Drosophila* Stock Center. *esg^ts^* refers to *tub-GAL8O^ts^, esg-GAL4, UAS-GFP* (II).

For tumor induction, virgin *esg^ts^* female flies were crossed with *UAS-yki^act^*, or *UAS-Ras1A*, or *w*^1118^ males. Crosses were kept at 18°C with regular flipping of the vials every 4–6 days. For the experiment, 15 females and 2 males were collected every 24–48 hours and incubated at 29°C to induce *yki^act^* expression in the gut.

For crosses involving *UAS-BirA*-ER* and *UAS-Yki^act^*, flies were maintained at 25°C. Once the F1 progeny were collected, 15 females and 2 males were transferred to food containing 100 µM biotin and incubated at 29°C for 8-12 days, with food being changed every two days, until sample collection or further processing.

### Knockdown and overexpression experiments

The following stocks were used for RNAi knockdown and overexpression experiments: Bloomington *Drosophila* Stock Center (BDSC)-BL58993, BL55276 (*UAS-egr RNAi*); BL32934, BL34938, BL27650 (*UAS-dl-RNAi*); BL29514 (*UAS-dif-RNAi*); BL55777 (*UAS-Flag-rel68*), BL31044, BL31477 (*UAS-Tl-RNAi*), BL33383 (*UAS-PGRP-LC-RNAi*); *UAS-Lsp1gamma-RNAi*; BL65086 (*UAS-Tg RNAi*); Vienna Drosophila Resource Center-330748 (*UAS-fon-RNAi*); 330200 (*UAS-FBP1 RNAi*); FlyORF Stocks-F000891, F001717 (*UAS-Sd-3XHA*), F000638 (*UAS-dl-3XHA*).

### Biotin food preparation and hemolymph collection

Biotin food was prepared by creating a stock solution (approximately 1 mM) by dissolving solid biotin (Sigma B4639) in water and adjusting the pH to ∼7.2. The stock solution was stored at -20°C. For fly food containing 50 µM biotin, standard fly food was liquefied using a microwave until a smooth consistency was achieved. Biotin was added at a final concentration of 100 µM, along with 5% yeast pellets (w/v). The mixture was thoroughly blended and poured into vials or bottles, which were then covered with cheesecloth and left to dry overnight. Once dried, the vials were capped and stored at 4°C until needed.

### Hemolymph isolation from adult flies

To collect hemolymph, 20-30 microcentrifuge tubes (0.5 mL) were prepared by piercing a hole in the bottom using a 25-gauge needle. An equal number of 1.5 mL microcentrifuge tubes were prepared by adding 15 µL of phosphate-buffered saline (PBS) to each. On a CO2 pad, 25 flies from each genotype were punctured in the abdomen using a tungsten needle (Fine Science Tools). The punctured flies were then transferred to the pierced 0.5 mL tubes, which were placed inside the 1.5 mL tubes containing PBS. The tube assembly was centrifuged at 2348 × g for 5 minutes at 4°C to collect the hemolymph. After removing the 0.5 mL tube, the hemolymph in PBS was centrifuged in the 1.5 mL tube at 2348 × g for 5 minutes at 4°C. The supernatant was transferred to another 1.5 mL tube and centrifuged at 14,000 × g for 15 minutes at 4°C. Finally, the supernatant was collected, flash-frozen, and stored at -80°C until further use.

### Immunoblotting and immunoprecipitation of biotinylated proteins

Hemolymph samples were pooled together. Protein concentration was calculated using a BCA kit (Pierce 23225). BCA normalized protein samples were mixed with an equal volume of 4x SDS sample buffer and boiled for 5 minutes at 95°C. 10 μg/sample was loaded onto a 4-20% Mini-PROTEAN TGX PAGE gel (Biorad 4561095), transferred to Immobilon-FL PVDF membrane (Millipore IPFL00010), incubated in PBS + 0.1% Tween (PBST) for 15 minutes, and blocked overnight in 3% BSA in PBST (PBST-BSA) at 4°C. To detect biotinylated proteins, blots were incubated with 0.3 μg/mL streptavidin-HRP (Thermo Fisher S911) in PBST-BSA for 1 hour at room temperature. Blots were washed four times with PBST and exposed using Pico Chemiluminescent Substrate (Thermo Fisher 34577).

Streptavidin magnetic beads (Pierce 88817) were washed (using a magnetic stand for separation) twice in 1X RIPA+1X Protease inhibitor (PI) buffer and resuspended in 1 X RIPA +PI buffer. For immunoprecipitation-immunoblot experiments 4 mg (3mg/ml) of protein was combined with 40□µL of beads. 10ul of input sample was used for immunoblots. Samples were incubated overnight at 4□°C with end-over-end mixing. On the next morning, beads were washed twice in 1X RIPA+PI buffer, once in 2□M urea in 10□mM Tris (pH□=□8.0). 30ul of immunoprecipitation sample was stored for immunoblot analysis and the rest of the sample was used for Mas Spec analysis. To elute bound biotinylated proteins, washed beads were resuspended in equal volumes of 1× protein loading buffer, boiled for 5□min, and vortexed. Biotinylated proteins in the input, flowthrough and immunoprecipitation samples were detected by immunoblot blot using streptavidin-HRP.

## Mass Spectrometry

### On-bead trypsin digestion

Samples were collected and enriched with streptavidin magnetic beads, washed twice with 200 μl of 50 mM Tris-HCl buffer (pH 7.5), transferred into new 1.5-ml Eppendorf tubes, and washed two more times with 200 μl of 50 mM Tris (pH 7.5) buffer. Samples were incubated in 0.4 μg of trypsin in 80 μl of 2 M urea/50 mM Tris buffer with 1 mM DTT for 1 h at room temperature while shaking at 1000 rpm. Following pre-digestion, 80 μl of each supernatant was transferred into new tubes. Beads were then incubated in 80 μl of the same digestion buffer for 30 min while shaking at 1000 rpm. Supernatant was transferred to the tube containing the previous elution. Beads were washed twice with 60 μl of 2 M urea/50 mM Tris buffer, and these washes were combined with the supernatant. The eluates were spun down at 5000 × *g* for 30 s, and supernatants transferred to a new tube. Samples were reduced with 4 mM DTT for 30 min at room temperature, with shaking. Following reduction, samples were alkylated with 10 mM iodoacetamide for 45 min in the dark at room temperature. An additional 0.5 μg of trypsin was added, and samples were digested overnight at room temperature while shaking at 700 rpm. Following overnight digestion, samples were acidified (pH□<□3) with neat formic acid (FA), to a final concentration of 1% FA. Samples were spun down and desalted on C18 StageTips as previously described. Eluted peptides were dried to completion and stored at □80°C.

### Tandem mass tag labeling and StageTip peptide fractionation

Desalted peptides were labeled with tandem mass tag (TMTPro) reagents (Thermo Fisher Scientific). Peptides were resuspended in 80 μl of 50 mM HEPES and labeled with 20 μl of 25 mg/ml TMTpro reagents in acetonitrile (ACN). Samples were incubated at room temperature for 1 h with shaking at 1000 rpm. TMT reaction was quenched with 4 μl of 5% hydroxylamine at room temperature for 15 min with shaking. TMT-labeled samples were combined, dried to completion, reconstituted in 100 μl of 0.1% FA, and desalted on StageTips.

The TMT-labeled peptide sample was fractionated by basic reverse phase fractionation. StageTips were packed with two disks of SDB-RPS (Empore) material and conditioned with 100 μl of 100% MeOH, followed by 100 μl of 50% MeCN/0.1% FA and two washes with 100 μl of 0.1% FA. Peptide samples were resuspended in 200 μl of 1% FA (pH□<□3) and loaded onto StageTips. Eight step-wise elutions were carried out in 100 μl of 20 mM ammonium formate buffer with 5%, 7.5%, 10%, 12.5%, 15%, 20%, 25% and 45% MeCN. Eluted fractions were dried to completion.

### LC–MS/MS

All peptide samples were separated with an online nanoflow Vanquish Neo UPLC liquid chromatography system (Thermo Fisher Scientific) and analyzed on an Orbitrap Exploris 480 mass spectrometer (Thermo Fisher Scientific). The LC system, column and platinum wire used to deliver electrospray source voltage were connected via a stainless-steel cross (360□mm, IDEX Health & Science, UH-906x). The column was heated to 50C using a column heater sleeve (Phoenix-ST). Each sample was injected onto an in-house packed 20□cm□×□75□μm internal diameter C18 silica picofrit capillary column (1.9□um ReproSil-Pur C18-AQ beads, Dr. Maisch GmbH; PicoFrit 10□μm tip opening, New Objective, PF360-75-10-N-5). Mobile phase flow rate was 200□nl□min^−1^, composed of 3% acetonitrile/0.1% formic acid (Solvent A) and 90% acetonitrile/0.1% formic acid (Solvent B). The 154-min LC–MS/MS method used the following gradient profile: (min:%B) 0:1.8;1:5.4; 129:27; 138:54; 139:81; 144:81; 144.1:45; 153:45. Data acquisition was done in the data-dependent mode acquiring HCD MS/MS scans (*r*□=□45,000) after each MS1 scan (*r*□=□60,000) on the top 20 most abundant ions using a normalized MS1 AGC target of 100% and an MS2 AGC target of 30%. The maximum ion time used for MS/MS scans was 105□ms; the HCD-normalized collision energy was set to 34; the dynamic exclusion time was set to 20□s, and the peptide match and isotope exclusion functions were enabled. Charge exclusion was enabled for charge states that were unassigned, 1 and more than 6.

### Analysis of Mass spectrometry data

MS data was processed using Spectrum Mill v.7.11 (proteomics.broadinstitute.org). For all samples, extraction of raw files retained spectra within a precursor mass range of 600–6000□Da and a minimum MS1 signal-to-noise ratio of 25. MS1 spectra within a retention time range of ±60□s, or within a precursor *m*/*z* tolerance of ±1.4□*m*/*z* were merged. MS/MS searching was performed against a *Drosophila* database containing 264 common laboratory contaminants. Digestion parameters were set to ‘trypsin allow P’ with an allowance of four missed cleavages. The MS/MS search included fixed modification of carbamidomethylation on cysteine. TMTpro16 was searched using the full-Lys function. Variable modifications were acetylation and oxidation of methionine. Restrictions for matching included a minimum matched peak intensity of 30% and a precursor and product mass tolerance of ±20□ppm.

Peptide spectrum matches were validated using a maximum false discovery rate threshold of 1.2% for precursor charges 2 to 6 within each LC–MS/MS run. The Protein FDR was set to 0%. TMTpro reporter ion intensities were corrected for isotopic impurities in the Spectrum Mill protein/peptide summary module using the afRICA correction method that implements determinant calculations according to Cramer’s Rule. Fully quantified proteins were used for the final dataset. We used the Proteomics Toolset for Integrative Data Analysis (Protigy, Broad Institute, https://github.com/broadinstitute/protigy) to calculate moderated *t*-test *P* values. P values were adjusted for multiple hypothesis testing using the Benjamini–Hochberg method. Median normalization was performed to center aggregate distribution of protein-level log2 ratios. Spectra were searched against the UniProt *Drosophila* database to generate the protein list. Statistical analysis and quality control were performed to filter the hits. Proteins with more than one unique peptide sequence were selected. Strong replicate correlation and clustering were observed by PCA confirming a good correlation between technical and biological replicates (Supplementary Figures 1a, 1b, 1c).

Protein hits were selected based on adjusted P value (<0.05) and the enrichment ratio of experimental sample versus control sample. The ratio threshold was customized for each comparison (example-Yki^act^vs control) and determined by 10% false positive rate (FPR), which was calculated by an established approach based on the distribution of positive control genes such as annotated secreted genes as well as the negative control genes such as annotated transcription factors^25^. GSEA analysis was done by PANGEA^29^ and the gene sets selected include GLAD gene group annotation, gene ontology annotation of biological process using GO hierarchy, and GO slim2 annotation of biological process. The volcano plot and scatter plot visualization were done using VolcaNoseR^83^.

### RNA isolation, semiquantitative and RT quantitative PCR

For RNA isolation, 10-12 adult gut tissues were homogenized in TRIzol reagent (Ambion), followed by vigorous mixing with chloroform (Fisher) to enable phase separation. The supernatant was collected and processed using Direct-zol RNA MicroPrep columns (Zymo Research) following the manufacturer’s protocol. The eluted RNA was treated with TURBO DNA-free Kit (Invitrogen) to remove any residual DNA. Reverse transcription was performed using the iScript cDNA Synthesis Kit (Bio-Rad). Quantitative real-time PCR (qRT-PCR) was carried out on a CFX96 Real-Time System (Bio-Rad) using iQ SYBR Green Supermix (Bio-Rad). qRT-PCR reaction volume used was 10 µl (2.95 µl Nuclease free water + 0.05 µl 25 µM Primer pair mix+ 5 µ 2x SYBR Green+ 2 µl cDNA). Relative mRNA levels were determined using the ΔΔCt method, with mRNA levels normalized to *RP49* for gut tissues. List of primers used for quantitative PCR can be found in Supplementary Table 2.

### Immunostainings

For immunostainings, we followed the protocol outlined in Singh et al. (2019) ^84^. Adult *Drosophila* guts were dissected in cold 1X PBS and fixed for 20 minutes in 4% paraformaldehyde. The tissues were then subjected to four washes in a washing solution (1X PBS, 0.2% Triton-X-100, and 0.1% bovine serum albumin [BSA]) for 10 minutes each. Tissues were then incubated in a blocking solution (1X PBS containing 0.1% BSA and 8% normal goat serum) for 60 minutes.

Primary antibodies were diluted in the blocking solution, and the guts were incubated overnight at 4°C. After four washes with the washing solution, secondary antibodies, diluted at 1:200, were added and incubated for 90 minutes at room temperature. Subsequently, four additional 15-minute washes were performed using the washing solution. DAPI (1 µg/ml) was added for 20 minutes after a PBS wash. Tissues were finally washed with cold PBS for 10 minutes and mounted using VECTASHIELD Antifade Mounting Medium with DAPI (Vector Laboratories). Images were captured using Zeiss LSM980/Airyscan confocal microscope.

The following secondary antibodies were used: Alexa Fluor 555-conjugated goat anti-mouse IgG (1:200), Alexa Fluor 488-conjugated goat anti-mouse IgG (1:200), and Alexa Fluor 555-conjugated goat anti-rabbit IgG (1:200). Fluorescent images were captured using a Carl Zeiss LSM780 confocal microscope. Primary antibodies: anti Cleaved-Dcp-1 (Asp216) Rabbit Ab (Cell Signaling) (1:100); Rabbit Anti-Relish (Raybiotech, Inc. Cat: #RB-14-004) (1:200); Rabbit anti-B-GAL (Abcam #4761), Rabbit anti-phospho-Histone 3 antibody (Cell Signaling).

### Protein, triglyceride and glucose colorimetric measurements

Protein, triglyceride, and glucose quantifications were performed by modifying a previously published protocol. For each cross, three biological replicates were collected at the designated time point, with eight flies homogenized in PBS for each replicate. The samples were placed in an Abgene 96-Well 1.2 mL Polypropylene Deepwell Storage Plate (Thermo Fisher Scientific—AB0564) containing 200 µL of ice-cold PBS with 0.05% Triton-X-100 (Sigma) and 1.0 mm Zirconium Oxide Beads (Next Advance Lab Products—ZROB10). Homogenization was carried out using a TissueLyser II (QIAGEN) for 2-3 cycles of 30 seconds each, at an oscillation frequency of 30 Hz/s. Plates were centrifuged for 1 minute at 3197 g to remove debris, and the homogenate was immediately used for glucose and protein quantification. For triglyceride measurements, an additional heat treatment step was performed at 75°C for 10 minutes.

Quantifications were performed using a SpectraMax Paradigm Multi-mode microplate reader (Molecular Devices). For protein measurements, 5 µL of the homogenate was diluted in 20 µL of PBS and mixed with 200 µL of the BCA Protein Assay Kit (Pierce BCA Protein Assay Kit), followed by incubation for 30 minutes at 37°C with gentle shaking in 96-well Microplates (Greiner Bio-One). Glucose quantification involved mixing 10 µL of homogenate with 100 µL of Infinity Glucose Hexokinase Reagent (Thermo Fisher Scientific—TR15421), incubating for 30 minutes at 37°C in 96-well Microplates UV-Star (Greiner Bio-One – 655801) with gentle shaking. For triglycerides, 20 µL of the homogenate was mixed with 150 µL of Triglycerides Reagent (Thermo Fisher Scientific—TR22421) and incubated for 10 minutes at 37°C in 96-well Microplates (Greiner Bio-One) with gentle shaking. Values were calculated from standard curves using five serial dilutions (1:1) of BSA (Pierce BCA Protein Assay Kit), glycerol standard solution (Sigma), or glucose standard solution (Sigma).

### Climbing assay

20 adult female flies were transferred to a test tube with marks at 10cm. For climbing assays, flies were tapped to the bottom of the tube and the number of flies able to climb above 10□cm was counted after 10□s. Three biological replicates were performed for each cross, and each replicate repeated three times. After the climbing tests, flies were transferred back to regular food-containing vials.

### Lifespan assay

For lifespan analysis, 15 females and 2 males were collected every 24–48 hours and incubated at 29°C. 6 vials were used for each cross. Food was changed every two day and the number of dead flies was counted.

### PO activity measurements

We adapted the protocol from Hsi et al. (2023)^46^, to measure phenoloxidase (PO) activity ex vivo using hemolymph. Individual flies were bled onto small 1 cm² squares of Whatman filter paper (Whatman 1001 – Grade 1). Immediately after bleeding, 20 µL of 20 mM L-DOPA in PBS was applied to each blot. To prevent evaporation, the samples were covered and incubated for 30 minutes at 25°C. The blots were quickly dried by microwaving for 10 seconds, then allowed to air-dry completely for 30 minutes. Once dry, each blot was sealed with Scotch tape and scanned using an Epson Perfection 4490 Photo Scanner. Image intensity was quantified using FIJI, and background intensity was subtracted from all samples. To measure PO activity of hemolymph *ex vivo*, individual flies were bled onto small squares of Whatman filter paper (1cm^2^; Whatman 1001 – Grade 1). Immediately following each bleeding, 20μL of 20mM L-DOPA in PBS was applied to each blot. Samples were then covered to prevent evaporation and incubated for 30 minutes at 25°C. The blots were rapidly dried by heating in a microwave for 10 seconds and then allowed to completely air-dry for 30 minutes. Following drying, each blot was sealed in clear Scotch tape and scanned using an Epson Perfection 4490 Photo Scanner. Intensity of each blot was quantified in FIJI, and the mean background intensity was subtracted from all samples.

### Wound healing assay

Flies were anesthetized with CO2, and their abdomens punctured using a blunt 0.005-inch diameter tungsten needle. Flies were then returned to vials, and two hours post-injury, melanized wounds on the thorax were counted using a dissecting microscope.

### Bead aggregation assay

The protocol was modified from Hsi et al.^46^. Dynabeads were first washed twice with 10X PBS, then blocked overnight at 4°C on a rotator in a 0.1% BSA-PBS solution. The beads were then washed three times with 0.1X PBS and reconstituted in Ringer’s-PTU buffer (130 mM NaCl, 5 mM KCl, 1.5 mM CaCl2 × 2H2O, 2 mM Na2HPO4, 0.37 mM KH2PO4, 0.01% PTU) at a final concentration of 50%. The blocked beads were stored for up to two weeks at 4°C. A 2 µL sample of hemolymph was then mixed with 2 µL of blocked beads in a well of a 15-well glass slide, and the mixture was pipetted up and down ten times. The slide was incubated in a humid chamber at 25°C for 30 minutes. After incubation, bead aggregates were revealed by mixing the solution for 30 seconds with a 10 µL micropipette tip.

### Reactive Oxygen Species measurement

The protocol was adapted from Owusu-Ansah et al. ^85^. Guts were dissected in Schneider’s medium. Dissolved 1 µL of DMSO (Thermo Scientific, D23107) in 1ml of Schneider’s medium to give a final concentration of approximately 30uM. The guts were incubated with the dye for 5-7 minutes in a dark chamber, on an orbital shaker at room temperature, followed by three 5-minutes washes in Schneider’s medium. After mounting in vectashield, images were captured immediately using Zeiss LSM980/Airyscan confocal microscope.

### Chromatin immunoprecipitation

Chromatin isolation was conducted as previously published protocol with modifications^84^. A total of 50-100 F1 flies from the crosses between driver line *tubulin-GAL4, tubulin-GAL80^ts^* with *UAS-HA-Sd* (for HA-Sd immunoprecipitation), *UAS-Flag-Rel*^68^ (for Flag-Rel immunoprecipitation), and *UAS-HA-dl* (for HA-dl immunoprecipitation) were kept at 29’C for 2 days. F1 from *tubulin-GAL4, tubulin-GAL80^ts^* crossed with *w*^1118^ flies were used as negative controls for immunoprecipitation. Flies were quickly washed with absolute ethanol followed by wash in 1× PBS containing a protease inhibitor cocktail (PIC). The samples were then fixed in 1.5% formaldehyde for 20 minutes at room temperature. Crosslinking was halted by incubation in 1X glycine solution for 5 minutes, followed by three washes in cold 1× PBS with PIC, spin at 4000 x g, 5 minutes at 4°C. The flies were resuspended in 400 µl nuclear lysis buffer [1% SDS, 10 mM EDTA, 50 mM Tris-HCl (pH 8) and 1× protease inhibitor] and homogenized followed by sonication using a Bioruptor (30 cycles, pulse on for 30 seconds, pulse off for 30 seconds). After sonication, the lysates were clarified by centrifugation at 16000 x g for 10 minutes at 4°C, and the supernatant was stored at -80°C for further use. For the immunoprecipitation (IP), 200 µL (equivalent to 5-10 µg of chromatin) of the digested, cross-linked chromatin was used per IP, with 30 µL of diluted chromatin reserved as the input sample. Immunoprecipitation was done using 500 µL of the diluted chromatin incubated with 30 µL of pre-washed HA beads (Thermo Scientific, 88836), 30 µL of pre-washed Flag beads (Sigma-Aldrich, M8823), Rabbit IgG (Cell Signaling Technologies, 2729) along with ChIP-Grade protein G magnetic Beads (Cell Signaling Technologies, 9006) served as a mock-IP negative control, while an antibody against Histone H3 (D2B12) XP Rabbit mAB (Cell Signaling Technologies, 4620) served as a positive control. Samples were rotated overnight at 4°C.

Chromatin was eluted by adding 150 µL of 1X ChIP Elution Buffer to each IP sample, followed by incubation at 65°C with gentle vortexing (1,200 rpm) for 30 minutes. This step was repeated and both samples were pooled together. The protein G magnetic beads were pelleted using a magnetic separation rack, and the cleared chromatin supernatant was carefully transferred to new tubes. To reverse the cross-links, 6 µL of 5M NaCl and 2 µL of Proteinase K were added to each sample (including the 2% input sample), followed by overnight incubation at 65°C. Thereafter, the DNA was purified using phenol/chloroform extraction and ethanol precipitation followed by purification using the quick spin column and resuspended in 30 µL of TE buffer for subsequent qPCR analysis. The fold enrichment method comparing to non-tagged sample was used for normalization. The ChIP signals are divided by the control-HA/FLAG signals, representing the ChIP signal as the fold increase in signal relative to the background signal. Calculations were done as following: First dCt was calculated using Ct IP-CT control, followed by Fold change calculation using 2^-dCt.^ List of primers used for chromatin immunoprecipitation can be found in Supplementary Table 3.

## Author Contributions

Conceptualization, experiments design and writing by A.S. and N.P.; Experiments performance, data collection and analysis by A.S, L. L. and R.F.L.; Bioinformatic analysis by Y.H.; Mass-spectrometry performed by H.W., C.X., N.U., and S.C. All authors have read and agreed to the published version of the manuscript.

This article is subject to HHMI’s Open Access to Publications policy. HHMI lab heads have previously granted a nonexclusive CC BY 4.0 license to the public and a sublicensable license to HHMI in their research articles. Pursuant to those licenses, the author-accepted manuscript of this article can be made freely available under a CC BY 4.0 license immediately upon publication.

## Data Availability Statement

The data presented in this study are available on request from the corresponding authors. The original mass spectra and the protein sequence database used for searches have been deposited in the public proteomics repository MassIVE (http://massive.ucsd.edu) and are accessible at ftp://MSV000096735@massive.ucsd.edu.

## Acknowledgments

We would like to thank Dr. Justin A. Bosch for providing valuable insights and guidance on the BirA* labeling experiments, Dr. Ying Liu for sharing his Yki tumor snRNAseq data, Drs. Stephanie Mohr and Afroditi Petsakou for comments on the manuscript.

