## Supplemental Tables and Figures for "Cell-death induced immune response and coagulopathy promote cachexia in *Drosophila*"

**Supplementary Table 1:** List of identified proteins secreted from BirA* labeled tumorous and control guts. Column A: Flybase gene ID; Column B: Gene name; Column C: Unique Protein sequence identified in mass-spectrometry Column D: Presence of a signal peptide; Column E: Log fold change in $Yki^{act}$ compared to BirA* control; Column F: Log fold change in $Ras^{act}$ compared to BirA*control; Column G: Log fold change in BirA* compared to control; Column H: Log fold change in $Yki^{act}$ compared to $Ras^{act}$ Column I : Expression enrichment of genes in enterocytes (ECs) of $Yki^{act}$ gut compared to control guts, derived from snRNAseq data[34]. Column J: Expression enrichment of genes in intestinal stem cells (ISCs) of $Yki^{act}$ gut compared to control guts, derived from snRNAseq data; Column K: Protein size; Column L: Human Ortholog; Column M: Diopt Score; Column N: differential Expression in snRNAseq.

| FBgn | Gene | UniProt | signalP | logFC Yki-BioID | logFC Ras1A-BioID | logFC BioID-WT | logFC Yki-Ras1A | EC (Yki/WT) | ISC (Yki/WT) | Protein size | Human ortholog(s) | DIOPT sco | snRNA Expression |
|---|---|---|---|---|---|---|---|---|---|---|---|---|---|
| FBgn0002564 | Lsp1gamma | P11997 | Yes | 3.53 | -0.65 | -0.05 | 4.17 | 2.57 | 162.67 | 772 aa | – | – | yes |
| FBgn0033942 | Cpr51A | Q7JZW0 | Yes | 3.3 | -0.1 | -0.27 | 3.4 | 11.14 | 130.17 | 144 aa | – | – | yes |
| FBgn0034511 | GNBP-like3 | A1ZBU5 | Yes | 2 | -0.3 | -0.1 | 2.3 | 6 | 31 | 152 aa | – | – | Yes |
| FBgn0032281 | CG17107 | Q9VKS2 | Yes | 2.83 | 0.66 | 0.14 | 2.17 | 12 | 4 | 96 aa | – | – | yes |
| FBgn0266420 | Ote | P20240 | No | 2.09 | 0.13 | -0.13 | 1.96 | 2.11 | 2.8 | 424 aa | OTOG; | 1 | Yes |
| FBgn0040321 | GNBP3 | Q9NHA8 | Yes | 2.45 | 0.6 | -0.39 | 1.85 | 2.88 | 4 | 490 aa | – | – | yes |
| FBgn0035806 | PGRP-SD | Q9VS97 | Yes | 2.17 | 0.31 | -0.21 | 1.85 | 15.67 | 160 | 186 aa | PGLYRP1; | 8 | Yes |
| FBgn0014865 | Mtk | Q24395 | Yes | 2.76 | 1.06 | 0.3 | 1.7 | 18 | 13 | 52 aa | – | – | Yes |
| FBgn0004045 | Yp1 | P02843 | Yes | 2.55 | 0.87 | -0.11 | 1.68 | | | 439 aa | LIPC;PNLIP;PNLIPR | 4 | yes |
| FBgn0266405 | CG45045 | A0A6H2EDS4 | Yes | 2.42 | 0.75 | 0.23 | 1.68 | 6 | 2 | 42 aa | | | No |
| FBgn0036022 | CG8329 | Q9VT23 | Yes | 1.09 | -0.58 | -0.2 | 1.67 | 1 | 0 | 259 aa | CTRB1;CTRB2;CTRL | 2 | No |
| FBgn0005391 | Yp2 | P02844 | Yes | 2.5 | 0.85 | 0.03 | 1.65 | | | 442 aa | LIPC;PNLIP;PNLIPR | 4 | – |
| FBgn0039685 | Obp99b | Q9VAI6 | Yes | 1.02 | -0.62 | -0.58 | 1.64 | 0 | 0 | 149 aa | | | No |
| FBgn0024293 | Spn43Ab | A1Z6V5 | Yes | 1.76 | 0.13 | -0.09 | 1.63 | 4.05 | 7.67 | 393 aa | SERPINI2;SERPINI1 | 5 | yes |
| FBgn0032213 | CG5390 | Q9VL01 | Yes | 1.38 | -0.25 | -0.63 | 1.63 | 1.81 | 2 | 206 aa | TPSG1;TPSD1;TPSI | 3 | yes |
| FBgn0261989 | CG42807 | Q7JWE2 | Yes | 1.14 | -0.48 | -0.34 | 1.62 | 20 | 9 | 278 aa | | | Yes |
| FBgn0038105 | yellow-f2 | Q9VG08 | Yes | 1.05 | -0.58 | -0.3 | 1.62 | 5 | 1 | 452 aa | RGN; | 1 | No |
| FBgn0040735 | BomT2 | A1ZB62 | Yes | 1.45 | -0.14 | -0.32 | 1.59 | 5.44 | 9.83 | 76 aa | – | – | Yes |
| FBgn0037906 | PGRP-LB | Q8INK6 | Yes | 1.25 | -0.34 | -0.11 | 1.59 | 2.97 | 24.73 | 215 aa | PGLYRP2; | 10 | yes |
| FBgn0030251 | CG2145 | Q9VZ49 | Yes | 1.84 | 0.26 | -0.32 | 1.58 | 8 | 9 | 592 aa | ENDOU; | 15 | Yes |
| FBgn0028526 | CG15293 | Q9V3Y7 | Yes | 1.79 | 0.33 | -0.44 | 1.46 | 6.67 | 37 | 333 aa | – | – | yes |
| FBgn0032836 | CG10680 | Q9VIQ5 | Yes | 2.13 | 0.75 | -0.5 | 1.39 | 8.08 | 10 | 251 aa | – | – | Yes |
| FBgn0038252 | BigH1 | Q8MSC4 | No | 1.18 | -0.2 | -0.18 | 1.38 | 0.68 | 0.9 | 353 aa | H1-8;H1-0; | 2 | no |
| FBgn0034407 | DptB | A1ZBF6 | Yes | 2.97 | 1.6 | -0.08 | 1.37 | 7 | 3 | 120 aa | | | No |
| FBgn0033367 | PPO2 | Q9V521 | No | 1.85 | 0.47 | -0.73 | 1.37 | 3 | 0 | 684 aa | | | No |
| FBgn0034468 | Obp56a | Q9V8Y2 | Yes | 1.51 | 0.17 | 0.39 | 1.33 | 1.7 | 7 | 139 aa | | | Yes |
| FBgn0037891 | CG5214 | Q9VGQ1 | No | 0.86 | -0.46 | 0.17 | 1.32 | 1.71 | 5.13 | 468 aa | DLST; | 16 | Yes |
| FBgn0032472 | CG9928 | Q9VK39 | Yes | 2.51 | 1.23 | -0.17 | 1.29 | 3.79 | 6.14 | 106 aa | – | – | yes |
| FBgn0038535 | alt | Q960Y8 | No | 1.05 | -0.21 | -0.03 | 1.26 | 2.7 | 5.42 | 874 aa | RRBP1; | 4 | Yes |
| FBgn0022959 | yps | Q95RE4 | No | 1.32 | 0.08 | -0.44 | 1.24 | 1.85 | 3.44 | 340 aa | YBX2; | 12 | yes |
| FBgn0031296 | gNacalpha | Q9VPV7 | No | 1.33 | 0.1 | -0.46 | 1.23 | 0 | 0 | 347 aa | NACA;NACA2; | 8 | no |
| FBgn0054051 | CG34051 | Q6IH48 | Yes | 1.31 | 0.08 | -0.56 | 1.23 | 0.6 | 2 | 222 aa | | | No |
| FBgn0034512 | Bbd | A1ZBU8 | Yes | 1.88 | 0.66 | -0.13 | 1.22 | 8.76 | 9.92 | 245 aa | – | – | Yes |
| FBgn0053470 | BaraA1 | C0HLZ9 | Yes | 2.15 | 0.94 | -0.36 | 1.21 | | | 257 aa | | | – |
| FBgn0260746 | Ect3 | Q9VGE7 | Yes | 1.05 | -0.15 | -0.51 | 1.2 | 2.27 | 8 | 637 aa | GLB1L; | 17 | Yes |
| FBgn0039308 | CG11889 | Q9VBU0 | No | 0.91 | -0.27 | -1.21 | 1.17 | | | 420 aa | | | – |
| FBgn0034470 | Obp56d | Q8SY61 | Yes | 1.18 | 0.03 | -0.16 | 1.15 | 1.7 | 1.6 | 131 aa | | | Yes |
| FBgn0010228 | HmgZ | Q06943 | No | 1.15 | 0 | -0.37 | 1.15 | 3.39 | 6.96 | 111 aa | HMGB1;HMGB2;HM( | 4 | yes |
| FBgn0029715 | CG11444 | Q9W4J4 | No | 0.83 | -0.32 | -0.26 | 1.15 | 2.22 | 4 | 215 aa | PDAP1; | 15 | yes |
| FBgn0040582 | BomBc3 | Q9VD48 | Yes | 2.65 | 1.53 | -0.63 | 1.12 | 12.86 | 31.5 | 98 aa | – | – | yes |
| FBgn0028543 | NimB2 | Q7KTA1 | Yes | 2.32 | 1.21 | -0.34 | 1.12 | 3.38 | 2.8 | 403 aa | MEGF11;MEGF10; | 3 | yes |
| FBgn0039678 | Obp99a | Q9VAJ4 | Yes | 0.82 | -0.3 | -0.08 | 1.12 | 0.63 | 0.83 | 142 aa | | | No |
| FBgn0039682 | Obp99c | Q9VAI9 | Yes | 1.82 | 0.71 | -0.4 | 1.11 | 2.82 | 3.43 | 151 aa | – | – | yes |
| FBgn0013763 | Idgf6 | Q23997 | Yes | 0.63 | -0.47 | 0.02 | 1.11 | 1.75 | 3.76 | 452 aa | CHI3L2;OVGP1;CHI | 5 | Yes |
| FBgn0030160 | CG9691 | Q9W306 | Yes | 2.05 | 0.95 | -0.32 | 1.1 | 4.31 | 12 | 121 aa | – | – | Yes |
| FBgn0004240 | DptA | P24492 | Yes | 2.78 | 1.7 | -0.12 | 1.09 | 6 | 2 | 106 aa | | | Yes |
| FBgn0051313 | CG31313 | Q8SZN1 | Yes | 2.06 | 0.98 | -0.6 | 1.08 | 7.14 | 3.75 | 124 aa | CST11;CST7;CST3;( | 2 | Yes |
| FBgn0262057 | Spn77Ba | Q0E8C8 | Yes | 0.79 | -0.29 | -0.53 | 1.08 | 4.25 | 4.5 | 450 aa | SERPINB10; | 6 | yes |
| FBgn0012042 | AttA | P45884 | Yes | 2.54 | 1.47 | -0.05 | 1.07 | 15 | 18 | 221 aa | – | – | yes |
| FBgn0011694 | EbpII | Q23982 | Yes | 0.93 | -0.12 | -0.22 | 1.05 | 3 | 3 | 66 aa | | | Yes |
| FBgn0011695 | EbpIII | Q9W1C9 | Yes | 1.79 | 0.77 | -0.54 | 1.03 | 2.71 | 4.03 | 126 aa | – | – | yes |
| FBgn0030262 | Vago | Q9VZ35 | Yes | 1.52 | 0.49 | -0.88 | 1.03 | 1.89 | 14 | 160 aa | – | – | yes |
| FBgn0031561 | IM33 | Q9VQT8 | Yes | 1.41 | 0.39 | -0.16 | 1.02 | 3.91 | 5.86 | 82 aa | EPPIN-WFDC6;EPPI | 6 | Yes |
| FBgn0041581 | AttB | Q9V751 | Yes | 2.41 | 1.41 | 0 | 1.01 | 6 | 9 | 218 aa | | | Yes |
| FBgn0051680 | CG31680 | Q8INU9 | Yes | 1.38 | 0.37 | -1.11 | 1.01 | 1 | 0 | 147 aa | | | No |
| FBgn0000261 | Cat | P17336 | No | 1.35 | 0.36 | -0.53 | 0.99 | 2.42 | 6.5 | 3218 aa | CAT; | 16 | Yes |
| FBgn0010225 | Gel | Q07171 | Yes | 1.71 | 0.74 | -0.18 | 0.98 | 2.94 | 3.56 | 798 aa | GSN; | 15 | Yes |
| FBgn0032638 | SPH93 | Q9VJD7 | Yes | 1.43 | 0.45 | -0.36 | 0.98 | 2 | 1 | 494 aa | PRSS21;TPSG1;TP: | 2 | No |
| FBgn0039031 | Gbp3 | Q9VCT2 | Yes | 1.2 | 0.21 | -0.22 | 0.98 | 2.71 | 3 | 159 aa | | | Yes |
| FBgn0041579 | AttC | Q95NH6 | Yes | 3.44 | 2.47 | -0.12 | 0.97 | 12 | 3 | 241 aa | | | Yes |
| FBgn0031313 | CG5080 | Q7K3E2 | Yes | 1.53 | 0.56 | -0.37 | 0.97 | 2.6 | 5.4 | 554 aa | – | – | Yes |
| FBgn0034331 | BomBc2 | A1ZB61 | Yes | 1.25 | 0.31 | -0.39 | 0.93 | 4.15 | 4.67 | 154 aa | – | – | yes |
| FBgn0021906 | RFeSP | Q9VQ29 | No | 0.62 | -0.31 | 0.2 | 0.93 | 2.4 | 3.96 | 230 aa | UQCRFS1; | 15 | Yes |
| FBgn0061515 | endos | Q9VUB8 | No | 0.75 | -0.16 | -0.1 | 0.91 | 3.06 | 4.69 | 119 aa | ARPP19;ENSA; | 14 | Yes |
| FBgn0027090 | GlnRS | Q9Y105 | No | 0.62 | -0.29 | -0.34 | 0.91 | 2.33 | 4.23 | 778 aa | QARS1; | 17 | Yes |
| FBgn0033451 | Marc | A1Z803 | No | 0.6 | -0.31 | -0.23 | 0.91 | 1.6 | 3.33 | 340 aa | MTARC2;MTARC1; | 15 | Yes |
| FBgn0043841 | vir-1 | Q0E8R1 | Yes | 1.16 | 0.27 | -0.63 | 0.89 | 3.24 | 4.4 | 423 aa | – | – | yes |
| FBgn0004687 | Mlc-c | P54357 | No | 1.89 | 1.01 | -0.57 | 0.88 | 3.59 | 6.68 | 153 aa | MYL6; | 15 | Yes |
| FBgn0024814 | Clc | Q9VWA1 | No | 1.11 | 0.24 | -0.27 | 0.87 | 3.21 | 7.57 | 219 aa | CLTA; | 16 | yes |
| FBgn0002645 | Map205 | P23226 | No | 1.31 | 0.46 | -0.11 | 0.85 | 5.38 | 11.33 | 1185 aa | – | – | Yes |
| FBgn0043578 | PGRP-SB1 | Q70PY2 | Yes | 1.17 | 0.33 | 0.01 | 0.85 | 3 | 4 | 190 aa | PGLYRP2; | 7 | Yes |
| FBgn0041182 | Tep2 | Q9NFV7 | Yes | 1.05 | 0.2 | -0.41 | 0.85 | 4.11 | 7.21 | 1420 aa | CD109; | 14 | Yes |
| FBgn0026441 | ear | A0A0B4KG69 | No | 2 | 1.18 | -0.32 | 0.82 | 2.82 | 8.56 | 931 aa | MLLT1;MLLT3; | 11 | Yes |
| FBgn0036106 | CG6409 | Q9VTC3 | Yes | 1.15 | 0.33 | -0.33 | 0.82 | 4.3 | 12.11 | 360 aa | – | – | yes |
| FBgn0259975 | Sfp87B | B4ZJA7 | Yes | 1.63 | 0.82 | -0.29 | 0.81 | 1 | 0 | 87 aa | | | No |
| FBgn0037468 | CG1943 | Q9VI56 | No | 1.6 | 0.79 | -0.55 | 0.81 | 5.24 | 5.9 | 118 aa | JPT2; | 5 | Yes |
| FBgn0033875 | CG6357 | A1Z9I0 | Yes | 1.07 | 0.27 | -0.43 | 0.81 | 5.81 | 10.72 | 439 aa | CTSS;CTSK; | 4 | yes |
| FBgn0064237 | Idgf5 | Q8T0R7 | Yes | 0.98 | 0.17 | 0.3 | 0.81 | 3 | 8 | 444 aa | CHI3L2;OVGP1;CHI | 4 | Yes |
| FBgn0034200 | Gbp2 | Q7K533 | Yes | 1.69 | 0.9 | -0.59 | 0.79 | 1.89 | 2.29 | 456 aa | QRICH2; | 2 | Yes |
| FBgn0034860 | CG9812 | Q9W1Q4 | Yes | 1.48 | 0.69 | -0.64 | 0.79 | 0.44 | 3.2 | 245 aa | – | – | Yes |
| FBgn0036928 | Tom20 | Q95RF6 | No | 0.64 | -0.15 | -0.27 | 0.79 | 1.95 | 4.91 | 171 aa | TOMM20; | 17 | Yes |
| FBgn0050197 | CG30197 | Q8MKJ5 | Yes | 1.27 | 0.49 | -0.89 | 0.78 | 1.65 | 15.5 | 113 aa | SLPI;WFDC5;WFDC | 1 | yes |
| FBgn0015299 | Ssb-c31a | Q9VLR5 | No | 0.62 | -0.15 | -0.19 | 0.77 | 1.21 | 2.36 | 110 aa | SUB1; | 12 | Yes |
| FBgn0036619 | Cpr72Ec | Q9VV46 | Yes | 1.15 | 0.39 | -0.58 | 0.76 | 1 | 0 | 429aa | RRS1;SNRNP70;RB | 1 | No |
| FBgn0038465 | Irc | Q9VEP8 | Yes | 0.71 | -0.05 | -0.41 | 0.76 | 3.32 | 4 | 697 aa | EPX;MPO;PXDN;LP | 4 | Yes |
| FBgn0024183 | vig | Q9V426 | No | 1.52 | 0.77 | -0.42 | 0.75 | 1.95 | 5.07 | 490 aa | SERBP1; | 13 | Yes |
| FBgn0037027 | HIPP1 | Q9VPB0 | No | 0.84 | 0.09 | -0.24 | 0.75 | 2.31 | 5.23 | 926 aa | AUH;ECHDC2;CDY1 | 3 | Yes |
| FBgn0033820 | CG4716 | Q7JRN6 | Yes | 1.66 | 0.93 | -0.61 | 0.73 | 3.33 | 4.49 | 223 aa | – | – | Yes |
| FBgn0030321 | CG1703 | Q9VYW4 | No | 1.03 | 0.31 | -0.22 | 0.72 | 1.92 | 7.88 | 901 aa | ABCF1; | 15 | Yes |
| FBgn0040968 | CG14933 | Q9VKE7 | Yes | 0.83 | 0.12 | -0.71 | 0.72 | | | 77 aa | SPINK7; | 4 | – |
| FBgn0003462 | Sod1 | P61851 | No | 1.12 | 0.41 | -0.54 | 0.71 | 2.15 | 3.98 | 153 aa | SOD1; | 14 | yes |
| FBgn0031560 | CG16713 | Q9VQT7 | Yes | 0.93 | 0.24 | -0.13 | 0.69 | 3 | 5.5 | 82 aa | EPPIN-WFDC6;EPPI | 6 | Yes |
| FBgn0037810 | sle | Q8INM3 | No | 0.98 | 0.3 | -0.21 | 0.68 | 1.02 | 7.85 | 1430 aa | – | – | Yes |
| FBgn0034692 | CG13502 | Q9W295 | No | 1.37 | 0.71 | -1.23 | 0.66 | 1 | 0 | 410 aa | ODAD4; | 9 | No |
| FBgn0033179 | p47 | Q7K3Z3 | No | 0.66 | 0 | -0.14 | 0.66 | 2.71 | 7.33 | 407 aa | NSFL1C; | 16 | Yes |
| FBgn0051777 | CG31777 | Q8IQ06 | Yes | 1.43 | 0.82 | -0.98 | 0.61 | 5.29 | 14.5 | 109 aa | PAPLN;TFPI2;COL6 | 1 | Yes |
| FBgn0264922 | Sumo | O97102 | No | 3.11 | 2.53 | 0.08 | 0.58 | 2.19 | 4.27 | 90 aa | SUMO3; | 15 | yes |
| FBgn0052521 | Mnr | Q9VRF7 | Yes | 1.24 | 0.66 | -0.8 | 0.58 | 4.43 | 4.6 | 388 aa | – | – | yes |
| FBgn0010774 | Ref1 | Q9V3E7 | No | 0.74 | 0.16 | -0.53 | 0.58 | 3.27 | 8.7 | 266 aa | ALYREF; | 15 | Yes |
| FBgn0002930 | nec | Q7JWX3 | Yes | 0.66 | 0.1 | -0.58 | 0.57 | 3.45 | 5.33 | 476 aa | SERPINI1;SERPINB | 8 | Yes |
| FBgn0036058 | CG6707 | Q9VT69 | No | 0.6 | 0.03 | -0.27 | 0.57 | 3.6 | 4.29 | 266 aa | PIP4P1; | 16 | Yes |
| FBgn0052185 | edin | Q8IQR7 | Yes | 2.16 | 1.6 | 0.85 | 0.56 | 8.5 | 23 | 115 aa | – | – | Yes |
| FBgn0028936 | NimB5 | Q9VJU6 | Yes | 1.41 | 0.85 | -0.55 | 0.56 | 2.5 | 0 | 315 aa | STAB1;STAB2;PEAI | 2 | No |
| FBgn0036035 | CG18178 | Q9VT38 | No | 0.71 | 0.16 | -0.37 | 0.54 | 2.38 | 8.67 | 374 aa | – | – | Yes |
| FBgn0033351 | AIMP1 | A1Z7K8 | No | 0.69 | 0.17 | -0.41 | 0.52 | 1.69 | 4.5 | 323 aa | AIMP1; | 15 | Yes |
| FBgn0264750 | CG44008 | M9PBA3 | Yes | 1.52 | 1.05 | -0.64 | 0.47 | 3.46 | 6.44 | 79 aa | SPINK9;SPINK7;SP | 5 | Yes |
| FBgn0263198 | Acn | Q9VJ12 | No | 0.94 | 0.47 | -0.56 | 0.47 | 2.43 | 4.97 | 739 aa | ACIN1; | 10 | Yes |
| FBgn0003721 | Tm1 | P49455 | No | 0.88 | 0.41 | -0.21 | 0.47 | | | 282 aa | TPM1; | 15 | – |
| FBgn0052190 | NUCB1 | Q9VVK7 | Yes | 1.71 | 1.25 | -0.14 | 0.46 | 3.05 | 10.94 | 569 aa | NUCB2; | 14 | Yes |
| FBgn0033631 | Sod3 | Q7JR71 | Yes | 1.57 | 1.11 | -0.42 | 0.46 | 1.06 | 3.07 | 243 aa | SOD1; | 8 | Yes |
| FBgn0000547 | ed | Q9VQW7 | Yes | 0.66 | 0.22 | 0.45 | 0.44 | | | 1332 aa | NPHS1; | 3 | – |
| FBgn0262103 | Sik3 | A1ZBC4 | No | 0.81 | 0.4 | -0.57 | 0.41 | 2.66 | 6.13 | 1471 aa | SIK3;MARK4; | 4 | Yes |
| FBgn0287873 | zip | Q99323 | No | 1.75 | 1.37 | -0.11 | 0.39 | | | 2024 aa | MYH10; | 15 | – |
| FBgn0038914 | fit | Q9VD66 | Yes | 1.15 | 0.76 | -0.05 | 0.39 | 0.89 | 0.64 | 121 aa | | | No |
| FBgn0027599 | CG2790 | Q9W0X8 | No | 0.67 | 0.29 | -0.21 | 0.39 | 1.89 | 6.15 | 540 aa | DNAJC21; | 16 | Yes |
| FBgn0000719 | fog | P40795 | Yes | 2.44 | 2.06 | -0.28 | 0.38 | 8.55 | 6.21 | 730 aa | NECAP2; | 1 | Yes |
| FBgn0026721 | fat-spondin | Q7KN04 | Yes | 0.73 | 0.35 | -0.27 | 0.38 | 2 | 3.5 | 763 aa | SPON1; | 12 | Yes |
| FBgn0050069 | Sdb | A1Z9M5 | No | 0.81 | 0.44 | -0.18 | 0.37 | 2.43 | 4.38 | 4012 aa | SAXO2;SAXO1;MU( | 1 | Yes |
| FBgn0261260 | mgl | A8JTM7 | No | 0.73 | 0.36 | -0.14 | 0.37 | | | 4769 aa | LRP2; | 14 | – |
| FBgn0086408 | stl | Q9W1Z6 | Yes | 1.07 | 0.73 | -0.22 | 0.34 | 3.04 | 24.33 | 1136 aa | ADAMTS6;ADAMTS | 4 | Yes |
| FBgn0039219 | CG13630 | Q9VC48 | No | 0.65 | 0.34 | -0.42 | 0.31 | 1.41 | 7.5 | 374 aa | METAP1; | 16 | Yes |
| FBgn0033265 | coil | A1Z7A8 | No | 0.93 | 0.64 | -0.26 | 0.29 | 2.52 | 5.18 | 633 aa | – | – | Yes |
| FBgn0010772 | Xe7 | Q7KTT0 | No | 0.75 | 0.49 | -0.31 | 0.26 | 3.4 | 4.77 | 783 aa | AKAP17A; | 13 | Yes |
| FBgn0035542 | TP53INP | M9NFM7 | No | 0.91 | 0.66 | -1.15 | 0.25 | | | 417 aa | TP53INP1; | 7 | – |
| FBgn0030054 | Caf1-180 | Q9W3D1 | No | 0.76 | 0.56 | -0.47 | 0.2 | 2 | 5 | 1183 aa | CHAF1A; | 11 | Yes |
| FBgn0034753 | CG2852 | Q9W227 | Yes | 1.53 | 1.35 | -0.22 | 0.18 | 1.92 | 5.62 | 205 aa | PPIB; | 15 | Yes |
| FBgn0010434 | cora | Q9V8R9 | No | 0.69 | 0.52 | -0.39 | 0.17 | 4.14 | 8.3 | 1696 aa | EPB41L3;EPB41L1: | 13 | Yes |
| FBgn0039869 | Tbca | Q9V9U7 | No | 1.45 | 1.28 | -0.65 | 0.16 | 2.73 | 5.26 | 110 aa | TBCA; | 17 | Yes |
| FBgn0045862 | btz | A0A0B4KH64 | No | 1.23 | 1.08 | -0.36 | 0.14 | 2.94 | 12.25 | 806 aa | CASC3; | 11 | Yes |
| FBgn0033814 | Qsox1 | Q7JQR3 | Yes | 0.64 | 0.5 | -0.18 | 0.14 | 6.29 | 4.62 | 637 aa | QSOX1;QSOX2; | 16 | Yes |
| FBgn0031381 | Npc2a | Q9VQ62 | Yes | 1.92 | 1.79 | -0.09 | 0.13 | 11.79 | 11.31 | 148 aa | NPC2; | 16 | Yes |
| FBgn0033788 | CG13323 | Q4V4H7 | Yes | 1.49 | 1.36 | 0.04 | 0.13 | 4.89 | 10.91 | 112 aa | – | – | Yes |
| FBgn0032243 | Klp31E | Q9VKW4 | No | 0.67 | 0.54 | -0.48 | 0.13 | 2.77 | 4.31 | 1048 aa | KIF21A; | 15 | Yes |
| FBgn0261641 | CG42724 | A0A0B4KFB8 | No | 0.7 | 0.6 | -0.14 | 0.1 | 2.57 | 4.46 | 1123 aa | TCERG1; | 13 | Yes |
| FBgn0032283 | CG7296 | Q8SXV4 | Yes | 0.92 | 0.84 | 0.03 | 0.08 | 4.33 | 2 | 161 aa | HNRNPA2B1;COL21 | 1 | Yes |
| FBgn0031217 | Creld | Q9VPJ0 | Yes | 0.64 | 0.56 | 0.35 | 0.08 | 1.47 | 5.24 | 374 aa | CRELD2; | 14 | Yes |
| FBgn0067903 | IM18 | P82701 | Yes | 1.19 | 1.12 | 0.97 | 0.07 | | | 71 aa | | | – |
| FBgn0036847 | CNPYb | Q9VVW7 | Yes | 1.71 | 1.65 | 0.4 | 0.06 | 1.82 | 22 | 221 aa | CNPY4; | 15 | Yes |
| FBgn0032773 | fon | Q9VIX7 | Yes | 1.3 | 1.25 | -0.59 | 0.05 | 7.67 | 15.67 | 574 aa | NUP98;NCOA5; | 1 | Yes |
| FBgn0085446 | CG34417 | M9PDW8 | No | 0.65 | 0.61 | -0.42 | 0.05 | | | 4925 aa | SMTN; | 7 | – |
| FBgn0036745 | CG7484 | Q9VVJ7 | Yes | 1.15 | 1.12 | -0.36 | 0.03 | 2.46 | 9.33 | 178 aa | SELENOF; | 15 | Yes |
| FBgn0025740 | PlexB | Q9V4A7 | No | 1.12 | 1.11 | -0.25 | 0.02 | 3.21 | 5.01 | 2051 aa | PLXNA1;PLXNA2; | 9 | Yes |
| FBgn0004656 | fs(1)h | P13709 | No | 0.64 | 0.64 | -0.17 | 0 | | | 2046 aa | BRD4;BRD3; | 11 | – |
| FBgn0033663 | ERp60 | Q3YMU0 | Yes | 1.16 | 1.17 | -0.1 | -0.01 | 2.28 | 8 | 489 aa | PDIA3; | 17 | Yes |
| FBgn0039261 | Ythdf | Q9VBZ5 | No | 0.53 | 0.61 | -0.36 | -0.08 | 2.23 | 7.74 | 700 aa | YTHDF1; | 13 | Yes |
| FBgn0030734 | ERp44 | Q95TL8 | Yes | 1.14 | 1.23 | -0.22 | -0.09 | 1.78 | 6.56 | 412 aa | ERP44; | 17 | Yes |
| FBgn0035917 | Zasp66 | Q8IQB4 | No | 0.79 | 0.88 | -0.52 | -0.09 | 3.23 | 5.59 | 326 aa | PDLIM7;LDB3;PDLII | 3 | Yes |
| FBgn0027588 | GCS2alpha | Q7KMM4 | Yes | 0.9 | 1 | -0.13 | -0.1 | 2.03 | 7.46 | 924 aa | GANAB; | 16 | Yes |
| FBgn0040308 | Prx4 | Q9V3Q4 | Yes | 0.63 | 0.73 | -0.15 | -0.1 | 1.54 | 22.75 | 242 aa | PRDX4; | 16 | Yes |
| FBgn0259984 | kuz | Q9VJW9 | Yes | 1.13 | 1.24 | -0.28 | -0.11 | | | 1238 aa | ADAM10; | 14 | – |
| FBgn0034923 | Upf3 | Q9W1H3 | No | 0.51 | 0.63 | -0.73 | -0.11 | 2.04 | 18 | 492 aa | UPF3A;UPF3B; | 14 | Yes |
| FBgn0004598 | Fur2 | P30432 | No | 0.64 | 0.76 | -0.17 | -0.12 | 3.14 | 4.53 | 1679 aa | PCSK5; | 12 | Yes |
| FBgn0030787 | CG9609 | Q9VXB4 | No | 0.71 | 0.85 | -0.63 | -0.14 | 1.27 | 2.8 | 403 aa | GTF3A; | 10 | Yes |
| FBgn0039229 | Saf-B | Q8IMV6 | No | 0.53 | 0.68 | -0.82 | -0.15 | 2.94 | 4.19 | 928 aa | SAFB;SLTM; | 13 | Yes |
| FBgn0259785 | pzg | Q9VP57 | No | 0.54 | 0.73 | -0.23 | -0.18 | 2.85 | 5.95 | 996 aa | ZNF462; | 2 | Yes |
| FBgn0031089 | aspr | Q9W5X1 | Yes | 0.46 | 0.65 | -0.57 | -0.18 | 3.33 | 8.67 | 440 aa | DNER;EDIL3; | 1 | Yes |
| FBgn0002868 | MtnA | P04357 | No | 1.45 | 1.64 | -1.27 | -0.19 | 3.68 | 6.34 | 43 aa | MT1B;MT1H;MT2A;M | 1 | Yes |
| FBgn0004647 | N | P07207 | No | 0.82 | 1.04 | 0 | -0.22 | 3.52 | 4.77 | 2693 aa | NOTCH1;NOTCH2;N | 12 | Yes |
| FBgn0037718 | P58IPK | Q9VHA8 | Yes | 0.44 | 0.68 | 0.38 | -0.24 | 0.98 | 6.43 | 498 aa | DNAJC3; | 15 | Yes |
| FBgn0002781 | mod(mdg4) | Q86B87 | No | 0.4 | 0.65 | -0.43 | -0.25 | 2.48 | 4.14 | 540 aa | BTBD18; | 3 | Yes |
| FBgn0046214 | vig2 | Q9VBX3 | No | 0.69 | 0.96 | -0.38 | -0.27 | 1.56 | 4.43 | 443 aa | SERBP1; | 13 | Yes |
| FBgn0038043 | CG17202 | Q9VG76 | No | 0.61 | 0.89 | -0.7 | -0.28 | 2.05 | 2.74 | 181 aa | MYCBP; | 7 | Yes |
| FBgn0038842 | hdly | A8JR58 | Yes | 2.31 | 2.6 | -0.42 | -0.3 | 34.64 | 28.85 | 377 aa | – | – | Yes |
| FBgn0058160 | CG40160 | Q95RS6 | Yes | 0.58 | 0.89 | -0.38 | -0.3 | 3.45 | 11.05 | 421 aa | TPSAB1; | 4 | Yes |
| FBgn0031030 | Tao | Q0KHQ5 | No | 0.34 | 0.64 | -0.61 | -0.3 | 2.87 | 4.31 | 1039 aa | TAOK1; | 15 | Yes |
| FBgn0053861 | His1:CG33861 | Q4AB54 | No | 0.35 | 0.67 | -0.07 | -0.32 | | | 256 aa | H1-4;H1-5;H1-2; | 8 | – |
| FBgn0053858 | His1:CG33858 | Q4AB54 | No | 0.35 | 0.67 | -0.07 | -0.32 | | | | H1-4;H1-5;H1-2; | 7 | – |
| FBgn0053855 | His1:CG33855 | Q4AB54 | No | 0.35 | 0.67 | -0.07 | -0.32 | | | | H1-4;H1-5;H1-2; | 8 | – |
| FBgn0260456 | CG4806 | Q9W124 | No | 0.93 | 1.28 | -0.87 | -0.35 | 1.43 | 4.94 | 657 aa | RBM28; | 14 | Yes |
| FBgn0043903 | dome | Q9VWE0 | Yes | 0.34 | 0.69 | -0.07 | -0.35 | 3.06 | 9.12 | 1282 aa | PTPRQ; | 2 | Yes |
| FBgn0037930 | CG14715 | Q9VGK3 | Yes | 1.05 | 1.41 | 0.17 | -0.36 | 2.02 | 7.46 | 138 aa | FKBP2; | 14 | Yes |
| FBgn0286818 | Pdi | P54399 | Yes | 1.08 | 1.46 | 0.01 | -0.37 | 2.42 | 7.55 | 496 aa | P4HB; | 17 | Yes |
| FBgn0031016 | kek5 | Q9VWI6 | Yes | 0.53 | 0.91 | -0.45 | -0.37 | | | 931 aa | SLITRK4;LRRC24; | 3 | – |
| FBgn0033526 | Caf1-105 | A1Z898 | No | 0.32 | 0.69 | -0.35 | -0.37 | 1.88 | 6.67 | 747 aa | CHAF1B; | 15 | Yes |
| FBgn0051908 | CG31908 | Q8T9H8 | No | 0.82 | 1.21 | -0.89 | -0.39 | 2.23 | 5 | 392 aa | SZRD1; | 8 | yes |
| FBgn0033741 | CG8545 | A1Z909 | No | 0.73 | 1.13 | -0.36 | -0.39 | 1.84 | 5.13 | 891 aa | NOP2; | 13 | Yes |
| FBgn0026199 | myo | Q9V4F4 | No | 1.1 | 1.51 | -0.03 | -0.41 | 2.56 | 4.11 | 598 aa | MSTN; | 10 | yes |
| FBgn0003567 | su(Hw) | P08970 | No | 0.22 | 0.63 | -0.07 | -0.41 | 2.66 | 5.79 | 941 aa | ZNF358;ZNF746;ZB | 2 | Yes |
| FBgn0261618 | larp | Q9VAW5 | No | 0.23 | 0.65 | -0.47 | -0.42 | 3.06 | 4.17 | 1673 aa | LARP1B; | 14 | Yes |

**Supplementary Table 2:** List of primers used for quantitative PCR.

| | **Forward** | **Reverse** |
|---|---|---|
| fondue | ACTCTGCTCTGGGAAAACTCG | TCTCAACGGCACCACCTAATC |
| tiggrin | TCTGTCAGGGCTACGAGACC | GAGTTGTGGCACTGTTTGTCC |
| Eig71eE | CTAACTGTGGTCTGCTTAGTGG | CAACGCTTTCTCAATTACCTCCA |
| Idgf3 | AGCCCTACAATTCTGCACCC | CTGCTCAAGGTCCAATCGCTT |
| Eig71Ee | CTAACTGTGGTCTGCTTAGTGG | CAACGCTTTCTCAATTACCTCCA |
| fbp1 | ATCGTGGCGGCATTGATAAGG | CGAAGGGTGTCAAAGTCCTG |
| hemomucin | AGGTCATCAAGCTAACGTCCA | TGTTGCCCTGCGTATCAAAGG |
| hemolectin | TGGTTATGGCGGGATAAAGACG | GTTGCCCTGACTTCCCTGG |
| GNBP-like3 | TCTGTTCCTAgtcgCAATTTCC | GGTGAGTTGACCTTGACGGT |
| PGRP-lf | CACCCAGTGGGAAGTACCC | GTTCGCATCCTTCGGTTGC |
| tep-3 | TTCCCGCCTTAAGAACTGACA | CCGTCTGAACCAAAACCGTA |
| Glutactin | CGGAGACCCGATATGCACAG | TACCCAAGAAAGCGTTCACTG |
| ppo1 | TTGGAACTGCCCGATTCCTTC | TTCAGATCCACGTCCTTAGAGAA |
| ppo2 | GAGGAGTCTTTTGTGGTGCAG | GGTGAAGGTTGATGCCCAGA |
| ppo3 | ATCTTCACCAAAATGCAGACCG | TCGAgtcgATAAAGCGATCCG |
| sp7 | GTTGTAGGAATCCCAACCAGA | CTCCATCGAGGCACTGTGAG |
| puc | TCCGGCGGTCTACGATATAGAAA | AGCAATAGATGCGGGAAAA |
| d-jun | ACCTGAACACATCCACCCC | ATCCGGTGAGTTGATGACCAG |
| lsp1gamma | GCCTGTGTGACTGCCTTTAG | AGAGGCTCATCAATACGGTGA |
| AttA | CACAACTGGCGGAACTTTGG | AAACATCCTTCACTCCGGGC |
| DptB | ATGCATTTCACCGCTAGTCT | TGCCAGTGGTTCAGGCTG |
| metchnikowin | ATGCAACTTAATCTTGGAGCGA | GACGGCCTCGTATCGAAAATG |

| | | |
|---|---|---|
| drosomycin | GATGCCGACTGTCTCTCTGG | GACAGGTCTCGTTGTCCCAG |
| attacin D | ATGGAATGTCAGGCTTCAGGA | CCTGGAGTGGAGGCGAATAC |
| diptericin A | TACCCACTCAATCTTCAGGGAG | TGGTCCACACCTTCTGGTGA |
| imd | TCAGCGACCCAAACTACAATTC | TTGTCTGGACGTTACTGAGAGT |
| AttB | GCAATGGAGCTGGTCTGGAT | CCGATTCCTGGGAAGTTGCT |
| Dorsal | ATGTTTCCGAACCAGAACAATGG | CCGTTGTAGTTGAGGCTCTGT |
| Drosomycin B | GATGCCGACTGTCTCTCTGG | GACAGGTCTCGTTGTCCCAG |
| Defensin | CTGCAGCATAGCCGCCAGA | GCCGCCTTTGAACCCCTTGG |
| Drosocin | TTTTCCTGCTGCTTGCTTGC | GGCAGCTTGAGTCAGGTGAT |
| AttC | CGCCACCCAGAATCTACAGG | CTTAGGTCCAATCGGGCATCG |
| GNBP-like | TCTGTTCCTAgtcgCAATTTCC | GGTGAGTTGACCTTGACGGT |
| Spaetzle | GCGATTCCTTTGCAGGAGC | AATTAACTGCCAGGTgtcgTC |
| Toll | ATCTGAAGCATCCTTCGgtcg | GTTAGCCTAAACGTGGGATTCTC |
| Pelle | TGCAGCAGAGCTACAACGAA | CAGGATATTgtcgTGCCGGA |
| Dif | GGAGCCGACAAGCAATATAATCC | GTAGTTGCACACTTCGATGGT |
| Cactus | ATGCCGAGCCCAACAAAAG | CGCTAGTGGCTAGTGAGGAC |
| relish | CTTCCCGGAGGTTACACTGTG | GTGGGCTGTCCAAGTTAGTTT |
| edin | CAAGTGGgtcgGGAGGCTA | TCCGATTGTAgtcgAAATTCCG |
| Bombc3 | TGGTGAATGGCGTCTGTCTG | ACCACATTACCATCGCCAGG |
| vago | AAGCGATTCCTTATCGACCCT | GATCCTCTCGCGTGAAGACTT |
| eiger | AGCTGATCCCCCTGGTTTTG | GCCAGATCGTTAGTGCGAGA |
| eiger | AGCGAGTCGTCGATAATCTCC | GCATTCTCGTACTCCTTTTGG |
| spz | GACACCTGGCAGTTAATTGTCA | CGAAGTCACAGGGTTGATCCG |
| PGRP-LC | AGGCCGTCACAGTTACAGTG | GTGGTGGCCAGTACGATACC |
| PGRP-SD | GACAGCATGGAAACTCCCTTG | GTTTTGCAGATTTTGCATGTGC |

| | | |
|---|---|---|
| PGRP-SA | ACGGGCATAGCCTTTATCGG | TAATCCTCGCTCAGCTCACC |
| IDGF5 | CAGAGGTTGGAAAACTGGTGT | GTTCAGCCAGCGACATTTGG |
| IDGF6 | ATTCCGCCAGTTTCGTCAAGG | CGTAGACCAGATAGTCGCAGAA |
| RP49 | ATCGGTTACGGATCGAACAA | GACAATCTCCTTGCGCTTCT |
| GAPDH | CCAATGTCTCCGTTGTGGA | TCGGTGTAGCCCAGGATT |

**Supplementary Table 3: List of primers used for chromatin immunoprecipitation.**

| | FORWARD | REVERSE |
|---|---|---|
| *RP49* | ATCGGTTACGGATCGAACAA | GACAATCTCCTTGCGCTTCT |
| *Act5c*_Promoter | GTGCAGATAGCAGTAAACGTAAGC | CCCCAACTACTCATTGTATGCC |
| **Sd_ChIP** | | |
| *egr*_Promoter | TTACACAAAGTAAACAGCGCAGGC | GAACTGAGAACGGGAGCG |
| *egr*_intron | GCAAACTCTGGCTAGGGTATCTC | GTATGTCCCACACAATAACCAC |
| *Pvf1*_intron | GCAACAACAACAAGACGGC | CTTTTCATCGCTCTCCCTCC |
| *Impl2*_intron | CGAGCATTCGCGCACTTTAGAG | GGGGAAAACGCAACAGGTTGC |
| *upd3*_promoter | GCGCGCGGTGAAATTTAGAC | CTCGCAGACACGGCGATTC |
| **Rel_ChIP** | | |
| *Pvf1*_promoter | GATACCTGAGGCATGGATATATGC | CTTTGGCGAAATCTGATCAGC |
| *Impl2*_Intron | CGACGTTGGCAGAGAAAGAG | GGAGAGTCGCGAGTTCAAATG |
| *Impl2*_promoter | CTTTGTTAGCGCTGAGGAGCC | CAATCTGTAGGTGGCGCGT |
| *upd3*_promoter | CTGTCTTATCACCGCCTAGTCAAG | GCACTCGAAATGAGTGAGCATAG |
| **dl_ChIP** | | |
| *Pvf1*_promoter | GCTCTCCCAGCGTTAACTGTTAAC | CATTGCGCTCTCCGCTCTGTA |

| | | |
|---|---|---|
| *Impl2* | GTGCGTGTGTATGTGTGAGTG | GCTTCCCATCTGTGTGAGTATG |
| *upd3*_upstream | CGCTCAGCTGTGCTTTTTTATG | GAAGCGGAAGAGAGACAGGGAG |
| *upd3*_upstream | CGCTCAGCTGTGCTTTTTTATG | CCGCTCGAATCGCCAAAG |

**Supplementary Figure 1**

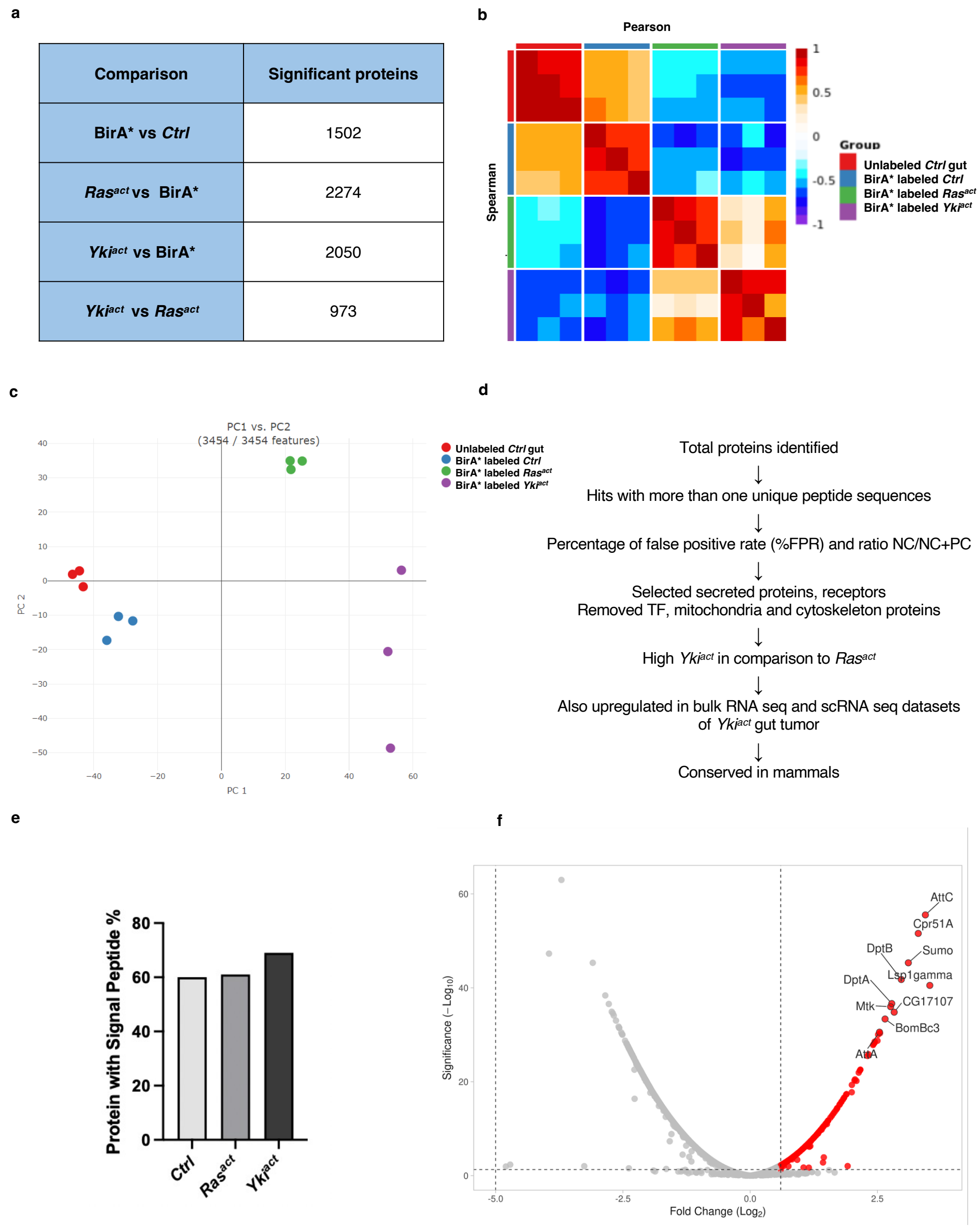


**Supplementary Figure 1. Identification and analysis of Yki$^{act}$ gut-derived proteins in hemolymph of adult flies.**

(a) Number of significant proteins identified in each sample (adjusted p-value < 0.05).

(b) Triplicates of the four samples analyzed for MS showing good Spearman and Pearson correlation.

(c) Principal component analysis showing good correlation among different replicates.

(d) Filters applied to narrow down the list of proteins identified by MS.

(e) Percentage of identified hits with signal peptide sequence in each sample. Cntrl is *EGT>BirA*, Ras$^{act}$* is *EGT>Ras$^{act}$ +BirA**, and *Yki$^{act}$ is EGT>Yki$^{act}$+BirA**

(f) Volcano plot to show top 10 hits identified in Yki$^{act}$ +BirA* gut secretomes compared to BirA* only.

**Supplementary Figure 2**

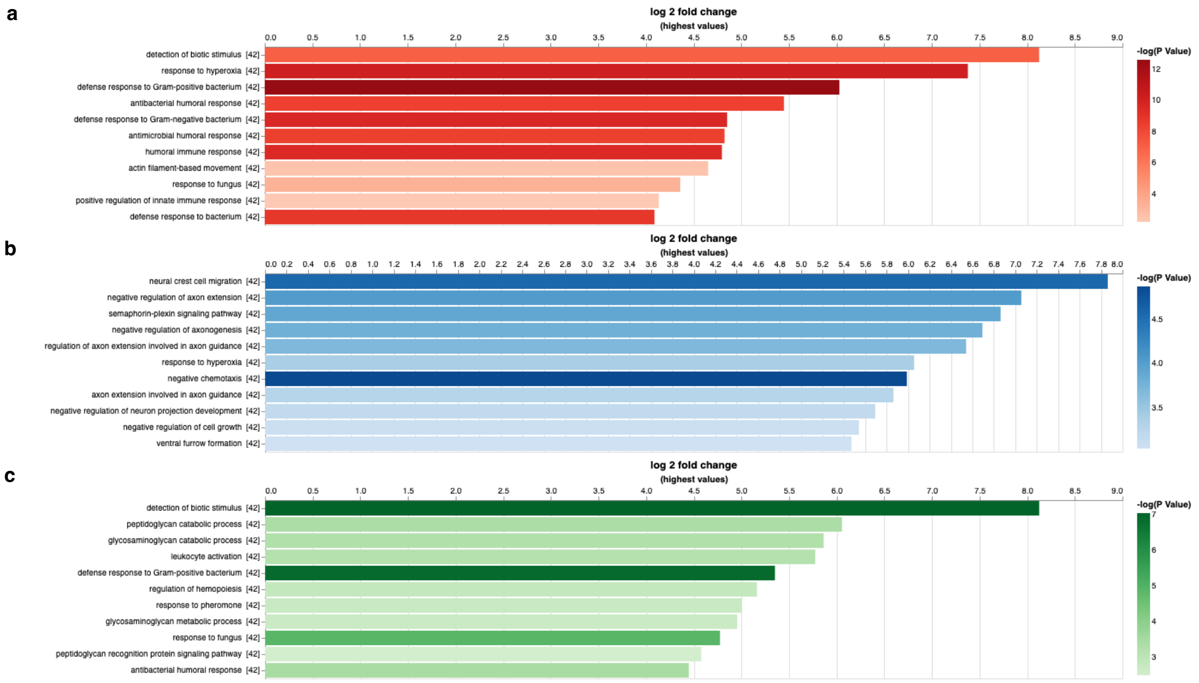


**Supplementary Figure 2.** Pathway enrichment analysis of top 100 proteins identified in Yki$^{act}$ (a), Ras$^{act}$ (b) and in both Yki$^{act}$ and Ras$^{act}$ (c) secretomes.

Supplementary Figure 3

a. modERN analysis

| TF | Gene | Peak Count | Location |
|---|---|---|---|
| Rel | *upd3* | 2 | Upstream |
| Rel | *upd3* | 1 | Intron |
| dl | *upd3* | 1 | Upstream |
| dl | *upd3* | | Intron |
| Dif | *upd3* | 2 | Upstream |
| Rel | *Impl2* | 3 | Upstream |
| Rel | *Impl2* | 2 | Intron |
| Rel | *Impl1* | | Upstream |
| dl | *Impl2* | 1 | Intron |
| Dif | *Impl2* | 1 | Upstream |
| Dif | *Impl2* | 1 | Intron |
| Rel | *Pvf1* | 1 | Intron |
| dl | *Pvf1* | 1 | Upstream |
| dl | *Pvf2* | 1 | Upstream |
| Dif | *Pvf1* | 1 | Upstream |
| Dif | *Pvf2* | 1 | Upstream |

b. AlphaFold 3 analysis

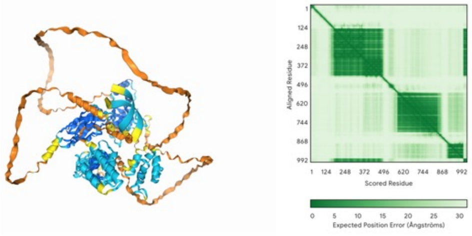


**(i) Rel-*upd3*  ipTM=0.92  pRM=0.41**

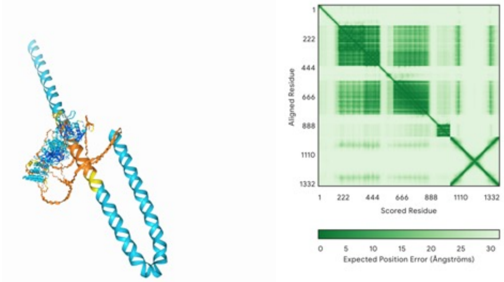


**(ii) Rel-*Impl2*  ipTM=0.26  pRM=0.48**

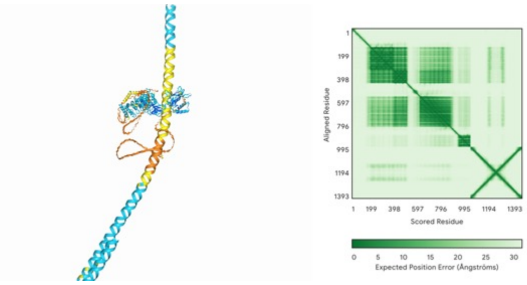


**(iv) Rel-*pvf1*  ipTM=0.22  pRM=0.42**

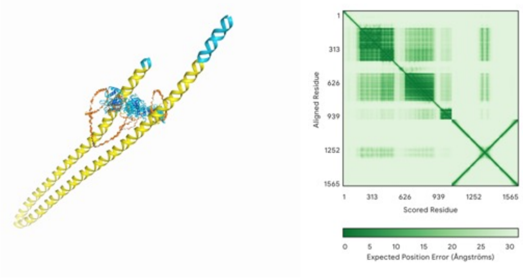


**(iii) Rel-*Impl2*_prom ipTM=0.24  pRM=0.39**

**Supplementary Figure 3: Analysis of NFκb and Sd transcription factor binding sites**

(a-b) NFκb transcription factor binding: (a) modERN analysis shows binding of NFκB transcription factors—Rel, dl, and Dif—to the regulatory sequences of *Pvf1*, *Impl2*, and *upd3*. (b) AlphaFold 3 analysis predicts binding of Rel to the regulatory regions of *upd3* (i), *Impl2* (ii, iii), and *pvf1* (iv). (c-d) Sd binding on *egr*: (c) modERN analysis suggests Sd binding to the regulatory sequences of *egr*. (d) AlphaFold 3 analysis indicating potential Sd binding to the *egr*.

**Supplementary Figure 4**

a

b

c

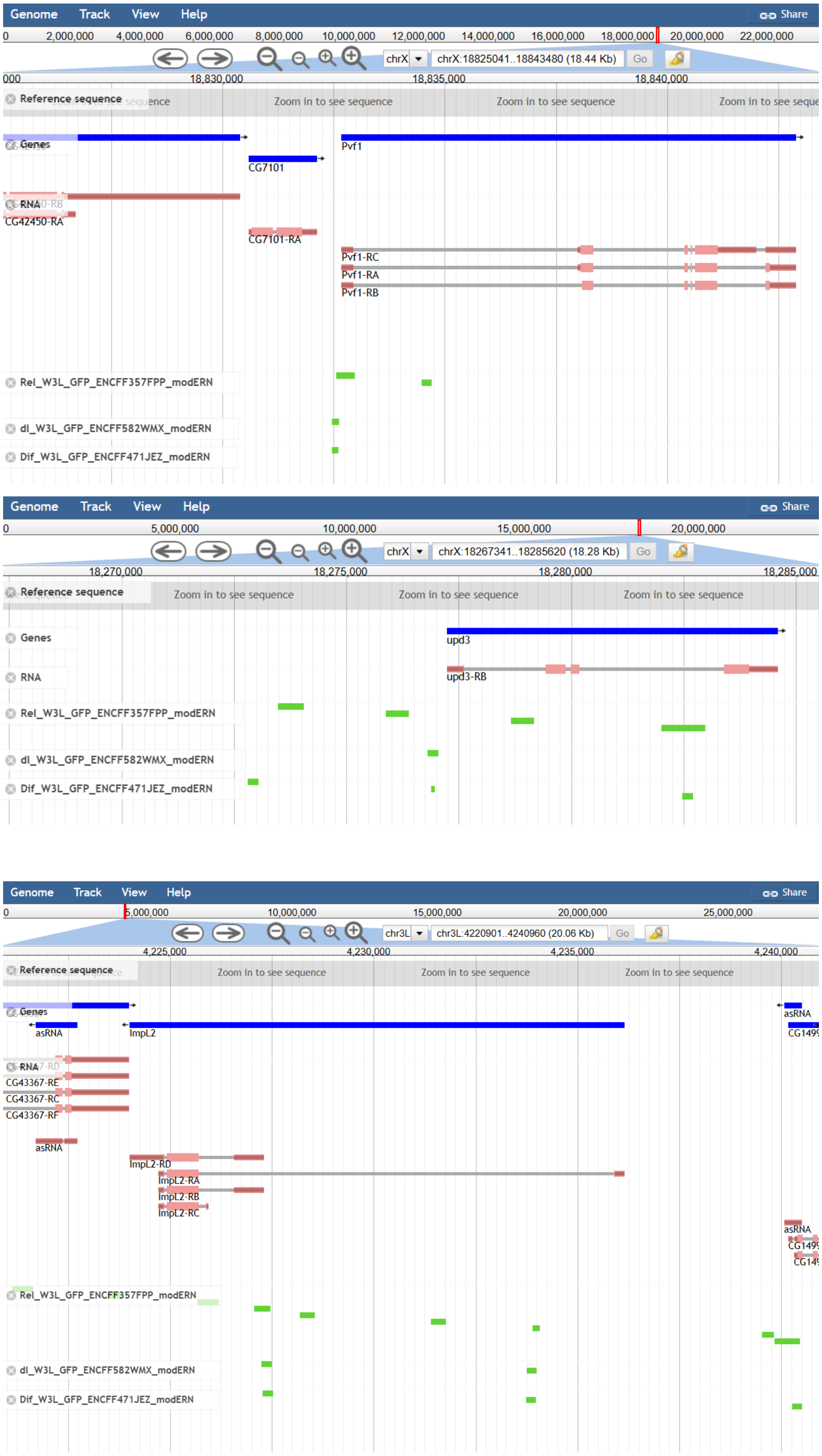

**Supplementary Figure 4: Bioinformatics analysis using Chip-seq datasets from modERN.** Chip-seq datasets for NFkB transcription factors from modERN near (a) *Pvf1*; (b) *upd3*; and (c) *ImpL2*.

**Supplementary Figure 5**

a

| TF | Gene | Peak Count | Location |
|---|---|---|---|
| Sd | *Eiger* | 4 | Intron |
| Sd | *Eiger* | 1 | Upstream |

b

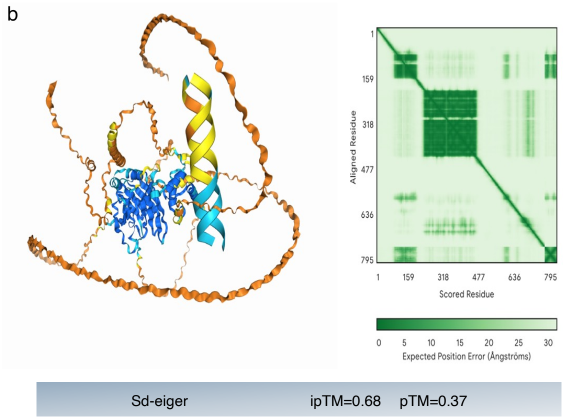


c

Gene Expression Counts

Gene: egr

| Sample Name | 00_indirect_flight_muscle | 01_follicle_cell | 07_enterocyte | 08_intestinal_stem_cell |
|---|---|---|---|---|
| 8-day WT | 13 | 99 | 56 | 60 |
| 8-day Yki | 65 | 89 | 236 | 223 |

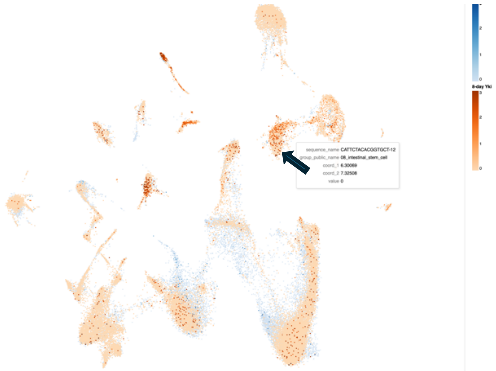

d

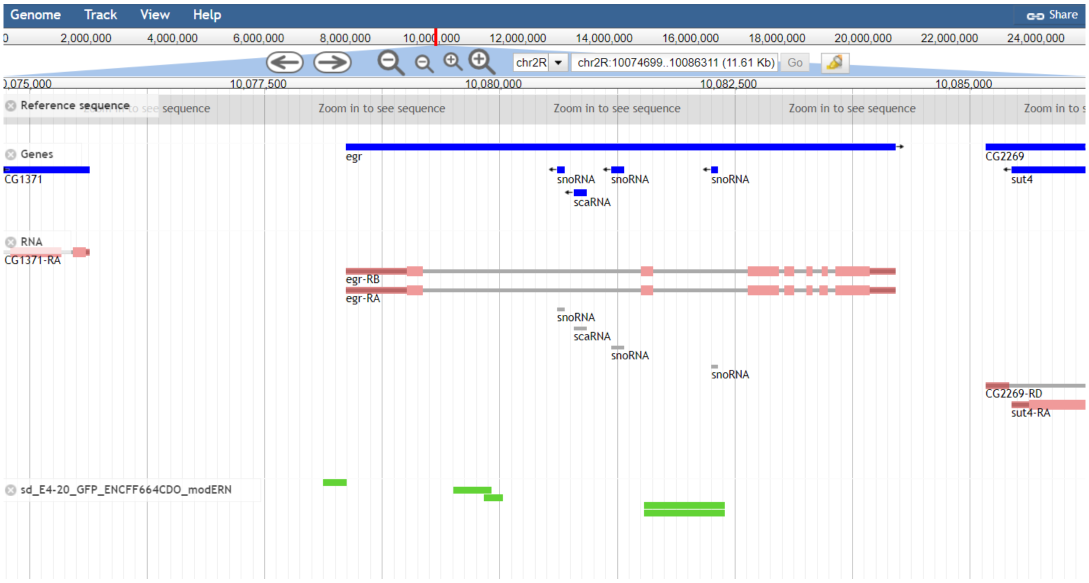

**Supplementary Figure 5: Analysis of Sd transcription factor binding sites on *egr* regulatory sequence.** (a) modERN analysis suggests Sd binding to the regulatory sequences of *egr*. (b) AlphaFold 3 analysis indicating potential Sd binding to the regulatory regions of *egr*. (c) snRNAseq analysis from $Yki^{act}$-whole body snRNAseq data[34] shows enrichment of *egr* gene expression in ISCs and EBs. (d) Bioinformatics analysis using Chip-seq datasets from modERN for Sd indicate peak signals near *egr*.
